# EMIn-depth profiling of calcite precipitation by environmental bacteria reveals fundamental mechanistic differences with relevance to application

**DOI:** 10.1101/850883

**Authors:** Bianca J. Reeksting, Timothy D. Hoffmann, Linzhen Tan, Kevin Paine, Susanne Gebhard

**Affiliations:** Department of Biology and Biochemistry, Milner Centre for Evolution, University of Bath, Claverton Down, Bath, BA2 7AY, United Kingdom; BRE Centre for Innovative Construction Materials, University of Bath, Claverton Down, Bath, BA2 7AY, United Kingdom

**Keywords:** ureolysis, self-healing concrete, microbial-induced calcite precipitation, *Bacillus*

## Abstract

Microbial-induced calcite precipitation (MICP) has not only helped to shape our planet’s geological features, but is also a promising technology to address environmental concerns in civil engineering applications. However, limited understanding of the biomineralization capacity of environmental bacteria impedes application. We therefore surveyed the environment for different mechanisms of precipitation across bacteria. The most fundamental difference was ureolytic ability, where urease-positive bacteria caused rapid, widespread increases in pH, while non-ureolytic strains produced such changes slowly and locally. These pH shifts correlated well with patterns of precipitation on solid media. Strikingly, while both mechanisms led to high levels of precipitation, we observed clear differences in the precipitate. Ureolytic bacteria produced homogenous, inorganic fine crystals, whereas the crystals of non-ureolytic strains were larger with a mixed organic/inorganic composition. When representative strains were tested in application for crack healing in cement mortars, non-ureolytic bacteria gave robust results, while ureolytic strains showed more variation. This may be explained by our observation that urease activity varied between growth conditions, or by the different nature and therefore material performance of the precipitate. Our results shed light on the breadth of biomineralization activity among environmental bacteria, an important step towards the rational design of bacteria-based engineering solutions.

## INTRODUCTION

Environmental protests worldwide have highlighted a growing demand for action on the threat of climate change. Human activity has greatly contributed to global warming, currently causing an estimated 0.2°C increase per decade due to past and ongoing emissions (Intergovernmental panel on climate change (IPCC) 2018 report). There is a clear need to reduce our carbon dioxide (CO_2_) emissions as well as to find novel ways in which we can consume CO_2_ as a means of decreasing overall levels. Geological sequestration of CO_2_ from large point source emitters into carbonate minerals, such as dolomite and limestone, is an emerging technology that could mitigate increasing CO_2_ concentrations (Mitchell *et al*., 2010). Microorganisms can act as biomediators to enhance carbon capture (Mitchell *et al*., 2010) by microbial induced calcite precipitation (MICP). This process leads to precipitation of CO_2_ as calcium carbonate (CaCO_3_) as a result of microbial metabolism and has been occurring on a geological scale for more than 2.5 billion years (Altermann *et al*., 2006). The formation by microbes of mineral carbonate structures, termed microbialites, is mediated by a large diversity of microorganisms, often working together in a consortium (Wilmeth *et al*., 2018). Indeed, cyanobacteria in association with heterotrophic bacteria are thought to be the principal contributors to the production of carbonate rocks during almost 70% of Earth’s history (3.5-0.5 Ga) (Altermann *et al*., 2006).

The success of MICP in biological engineering of the environment has led to the investigation of its potential for exploitation in other areas. These areas include bioremediation of heavy-metal contaminated soils via immobilisation of metals in precipitates (Achal *et al*., 2011;; Ceci *et al*., 2018), or wastewater treatment using this technology to remove excess calcium. Geotechnical engineering represents another area of interest, where MICP can improve soil properties by precipitating CaCO_3_ to bind sand particles together. Utilizing MICP in the construction industry is another rapidly emerging field. The production and maintenance of concrete bears heavy environmental and economic costs, with cement production contributing up to 5-8 % of global anthropogenic CO_2_ emissions (Huntzinger and Eatmon, 2009;; Souto-Martinez *et al*., 2018). Moreover, reinforced concrete structures face durability issues caused by cracking and subsequent ingress of water and ions, such as chlorides. This leads to corrosion of internal steel reinforcement and eventual structural failure. MICP can seal such cracks in concrete and reduce permeability to damaging substances, thereby extending the lifespan of structures and reducing the environmental burden caused by construction of replacement structures. This broad range of applications necessitates an understanding of how bacteria precipitate calcite and what the factors are that may limit their performance.

Most bacteria are capable of MICP under suitable conditions (Boquet *et al*., 1973), with key factors such as calcium concentration, concentration of dissolved inorganic carbon (DIC), pH, and availability of crystal nucleation sites affecting precipitation (Hammes and Verstraete, 2002). Bacterial metabolism results in an increase in both pH and DIC (through aerobic or anaerobic oxidation of organic compounds), and changes in solution chemistry (Power *et al*., 2011). This can lead to the oversaturation of Ca^2+^ and CO_3_^2-^ ions to facilitate the formation of calcium carbonate precipitates (Power *et al*., 2011). The negatively charged cell surfaces of bacteria further promote precipitation by attracting calcium ions and acting as nucleation sites for crystal formation (Zhu *et al*., 2016).

To date, the majority of studies of MICP have focussed on ureolytic bacteria (Zhu *et al*., 2016). These are associated with high rates of CaCO_3_ precipitation (Mitchell *et al*., 2010), caused by hydrolysis of urea (CO(NH_2_)_2_) to ammonia (NH_3_) and carbonic acid (H_2_CO_3_) (Mitchell *et al*., 2010). The reaction of ammonia with water to ammonium (NH_4_^+^) and hydroxide ions (OH^−^) causes an upshift in pH, while carbonic acid dissociates to generate bicarbonate (HCO_3_^−^). In the presence of calcium, this leads to the formation of a calcium carbonate (CaCO_3_) precipitate (Equations 1 and 2).

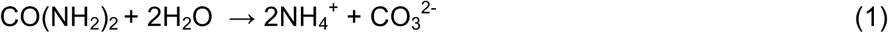

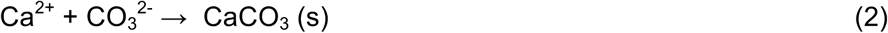

Ureolytic MICP has been used in a range of applications. Positive outcomes were obtained in the improvement of soil properties, however non-uniform rates of precipitation throughout the samples led to varying results (Zhao *et al*., 2014). In addition, the requirement for injection of the healing agent into the ground created problems where the rapid precipitation initiated by urease activity caused clogging at the injection site (Cheng and Cord-Ruwisch, 2014). In other applications, such as crack sealing in cementitious materials, ureolytic MICP has been shown to promote healing (Bang *et al*., 2010;; Wang *et al*., 2012). However, because of the dependency of the reaction on enzymatic activity, factors such as low temperature markedly decreased the efficiency of the reaction (Wang *et al*., 2017). In addition, ureolytic MICP requires the presence of urea, which may not always be feasible to provide, and the release of ammonia could contribute to environmental nitrogen loading (Jonkers *et al*., 2010).

To work around the challenges associated with ureolytic bacteria, the use of other metabolic pathways needs consideration. The high efficiency of ureolytic MICP has resulted in the assumption that non-ureolytic pathways cannot yield similarly high levels of precipitation and are thus an inferior strategy to achieve precipitation. This has limited our understanding of alternative pathways, and the factors affecting efficient calcite precipitation in non-ureolytic bacteria are not clear. Studies on self-healing concrete using alkaliphilic non-ureolytic bacteria, such as *Bacillus pseudofirmus* and *Bacillus cohnii*, have shown great promise and have led to similar levels of self-healing at the laboratory scale as ureolytic bacteria (Sharma *et al*., 2017;; Alazhari *et al*., 2018). To fully assess the potential of MICP for industrial and environmental use, detailed understanding of the microorganism, mechanism of precipitation, and application is therefore important.

Most studies of MICP have focussed on a small number of species and have thus restricted our understanding of the mechanisms of precipitation across bacteria. In addition, many previous studies have been primarily application-driven and have not always explored the deeper workings of how the bacteria precipitate minerals and what influences this ability. To build a strong basis on which to develop our understanding of MICP, we here took a broad view and surveyed in detail the ability of environmental bacteria to precipitate calcite. Focussing on bacteria that could grow in the alkaline conditions typical of high-calcium environments, we assessed the prevalence of MICP capability in environmental bacteria, their mechanisms of precipitation and resulting crystal morphologies as well as their suitability for application in self-healing concrete. Our study provides an understanding of the environmental potential for MICP, which is not only relevant to geomicrobiology but will facilitate a more design-based approach to industrial application of MICP.

## RESULTS AND DISCUSSION

### Calcite precipitation ability of environmental isolates

To gain a broader understanding of the capability of bacteria to precipitate calcium carbonate, we first performed a survey of a range of environments. In this, we focussed on sites expected to be calcium-rich, such as exposed limestone, caves and soils in areas with a limestone bedrock within the southwestern United Kingdom. Our isolation strategy targeted spore-forming bacteria, as their stress tolerance makes them more universally useful in potential applications. Moreover, as many of the high-calcium environments are in the alkaline pH range, we specifically sought for isolates that tolerate such conditions. Interestingly, the majority (89 %) of colonies obtained during primary isolations had visible crystal formation on the surface, indicative of biomineralization. This supports previous observations that this is a common trait among bacteria (Boquet *et al*., 1973). Of 74 isolates able to grow at pH 9, 31 displayed some growth at pH 10 on 0.25x B4 minimal medium and were chosen for further characterisation.

Preliminary identification based on partial sequencing of the 16S rDNA assigned all of the isolates into Bacillales (Fig. 1). The selection of spore-formers by heat-treatment can account for the prevalence of Bacillales, with most isolates grouping within the *Bacillus* genus. The predominant species represented were *Bacillus licheniformis* and *Bacillus muralis*. *Sporosarcina pasteurii* was included in the analyses as it represents the paradigm organism for industrial application of calcite precipitation. Interestingly, five of our isolates grouped within the same clade as *S. pasteurii* (Fig. 1).

**Figure 1.**
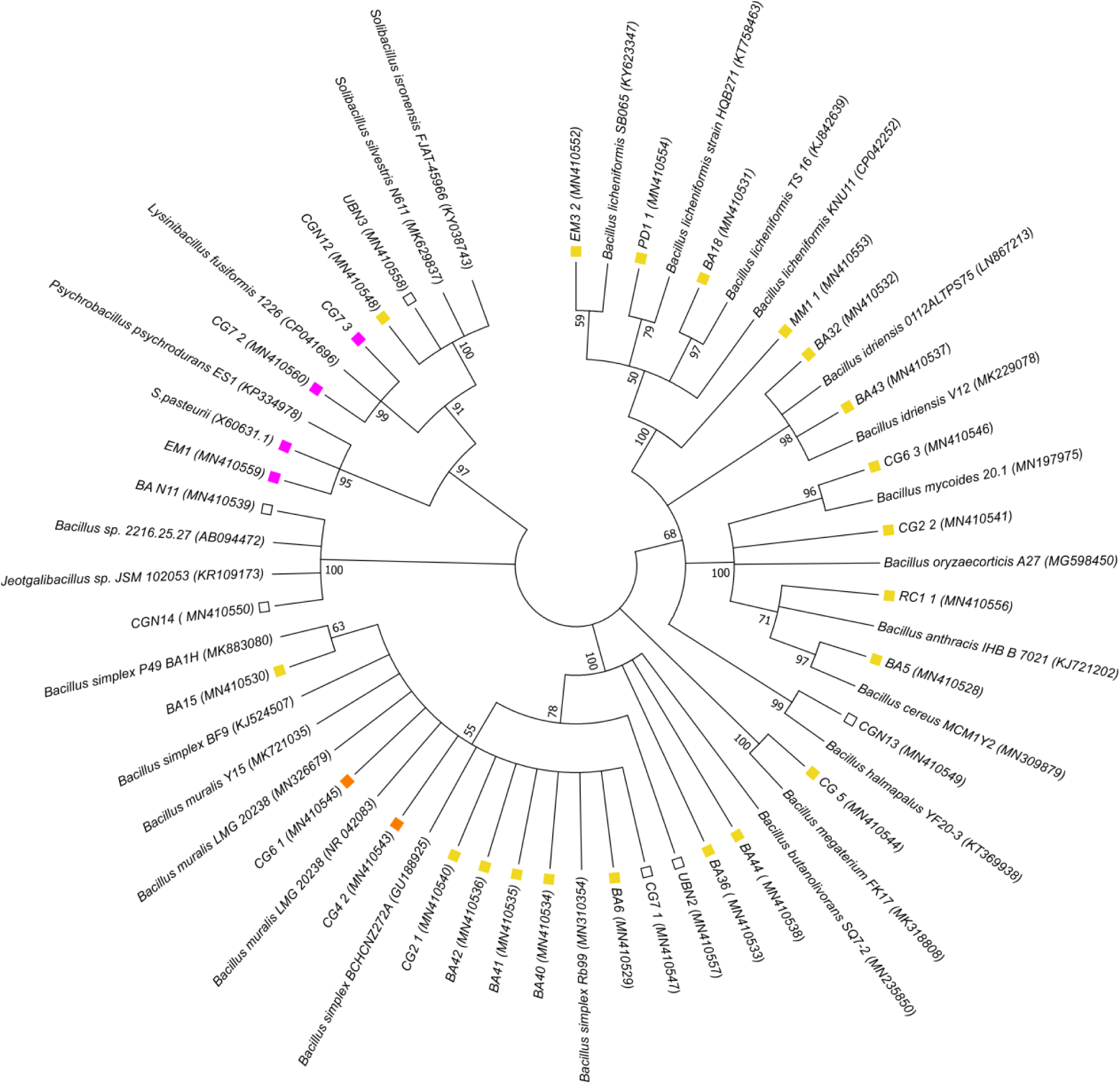
Phylogenetic analysis of selected calcite-precipitating isolates and their closest relatives. Yellow boxes, non-ureolytic isolates; orange boxes, isolates with inducible ureolytic activity; pink boxes, isolates that utilize urea as a primary nitrogen source; unfilled boxes, unknown ureolytic ability. The ureolytic potential of closest relatives is not shown. Accession numbers are indicated in parentheses. The percentage of trees in which associated taxa clustered together is shown next to the branches (1000 bootstraps).

### Mechanistic differences in calcite precipitation between isolates

The isolation strategy, with the exception of looking for alkali-tolerant spore formers, was unbiased for particular metabolic properties of the bacteria. A key feature of many known calcite precipitators is ureolysis, which leads to a rapid pH change in the environment due to the release of ammonia and subsequent production of OH^−^. We therefore tested each of our isolates for their ability to cleave urea, which is visible as a pink colour change in test broth containing urea and phenol red (Fig. 2 and Fig. S2). Five out of the 31 isolates taken forward for further characterisation were ureolytic. Two of these ureolytic isolates (CG4_2 and CG6_1) were most closely related to *B. muralis* whilst two others (CG7_2 and CG7_3) were identified as *Lysinibacillus fusiformis*. The final isolate (EM1) was identified as *Psychrobacillus psychrodurans* and was most closely related to *S. pasteurii*. With the exception of CG4_2 and CG6_1, the other ureolytic isolates formed a monophyletic clade with the strongly ureolytic *S. pasteurii* (Fig. 1).

**Figure 2.**
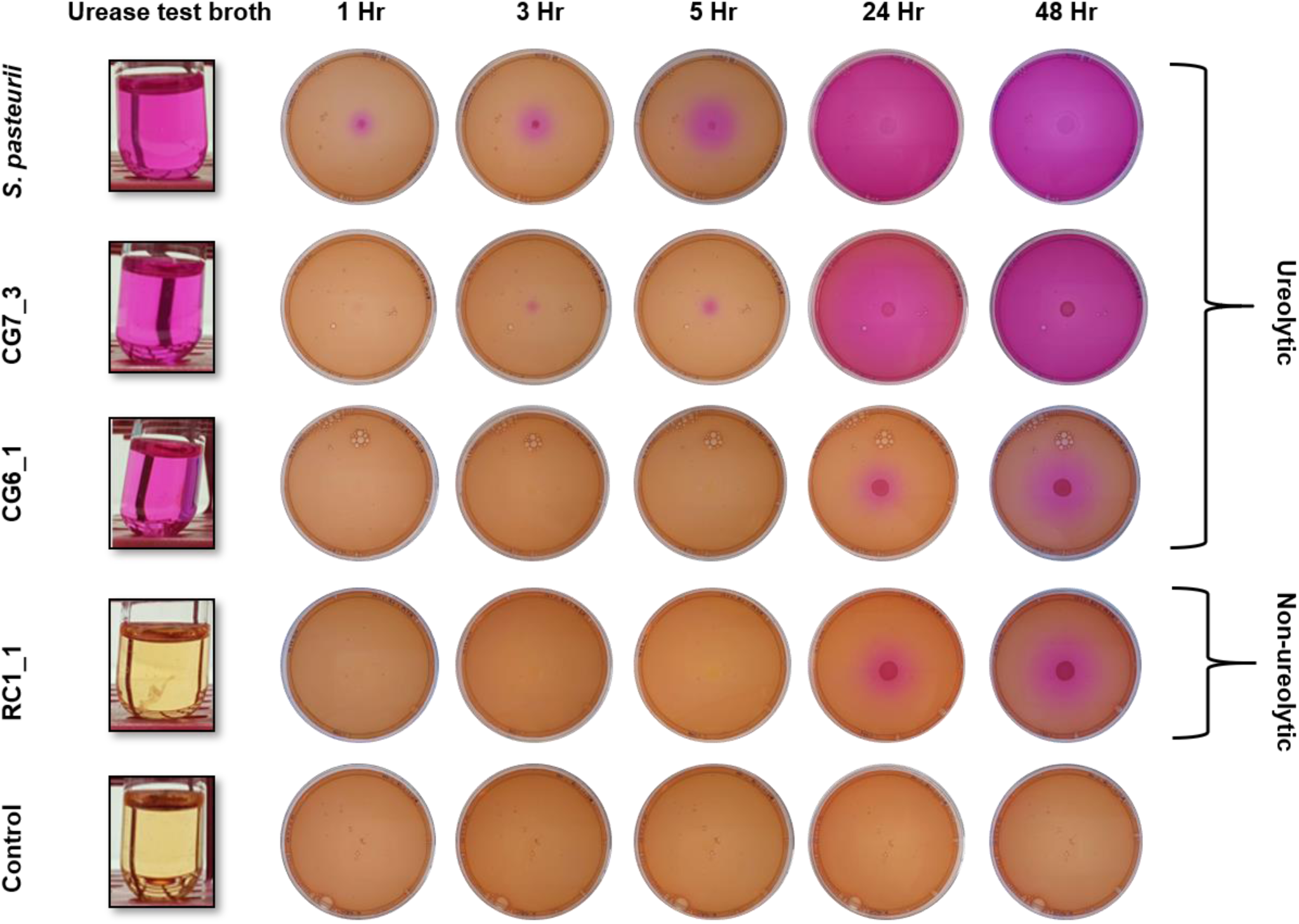
Diversity in ureolytic activity across environmental isolates. *Left*, urease test broth inoculated with the strains indicated to the left. *Right*, nutrient agar plates supplemented with phenol red, calcium acetate, and urea were inoculated in the centre with suspensions of the same strains. Pink shading indicates the pH change caused by ammonia release from ureolytic activity.

To determine the effects of ureolysis and the associated pH change on crystal formation, we repeated the experiment on solid media. Non-ureolytic strains showed only small changes in pH, as indicated by local colour changes in the immediate vicinity of the bacterial colony, due to general metabolic activity (Fig. 2, isolate RC1_1;; Fig. S2). In contrast, a rapid and wide-spread increase in pH was apparent in the model ureolytic bacterium *S. pasteurii*. Indeed, an increase in pH as indicated by a colour change from orange to pink was visible as early as 1 h after inoculation of the plate with *S. pasteurii*, and the pH of the entire plate was alkaline by 24 h (Fig. 2). Similarly, the closely related ureolytic isolate CG7_3 caused a rapid increase in pH, although the rate of pH change was slightly slower than seen in *S. pasteurii* (Fig. 2). The same behaviour was observed for isolate EM1 (Fig. S2).

We next investigated the patterns of calcium carbonate precipitation on the same agar plates and found that crystal formation correlated well with pH changes within the media. In non-ureolytic strains, crystals were visible only on the surface of the colony, corresponding to the observed localised pH change (Fig. 3A). In contrast, with ureolytic strains crystal precipitation reflected the global pH change and occurred throughout the media (Fig. 3B). The precipitation of calcium carbonate crystals throughout the media was likely due to a change in the saturation kinetics in the media, resulting from the rapid increase in pH caused by the urease-mediated release of ammonia from urea. Indeed, when urea was omitted for isolate CG7_3, only the localised pH changes associated with non-ureolytic isolates was seen (Fig. S3). In addition, crystals were only visible on the surface of the colony when urea was omitted (Fig. S4), indicating that there are mechanistic differences in calcite precipitation under ureolytic and non-ureolytic conditions.

**Figure 3.**
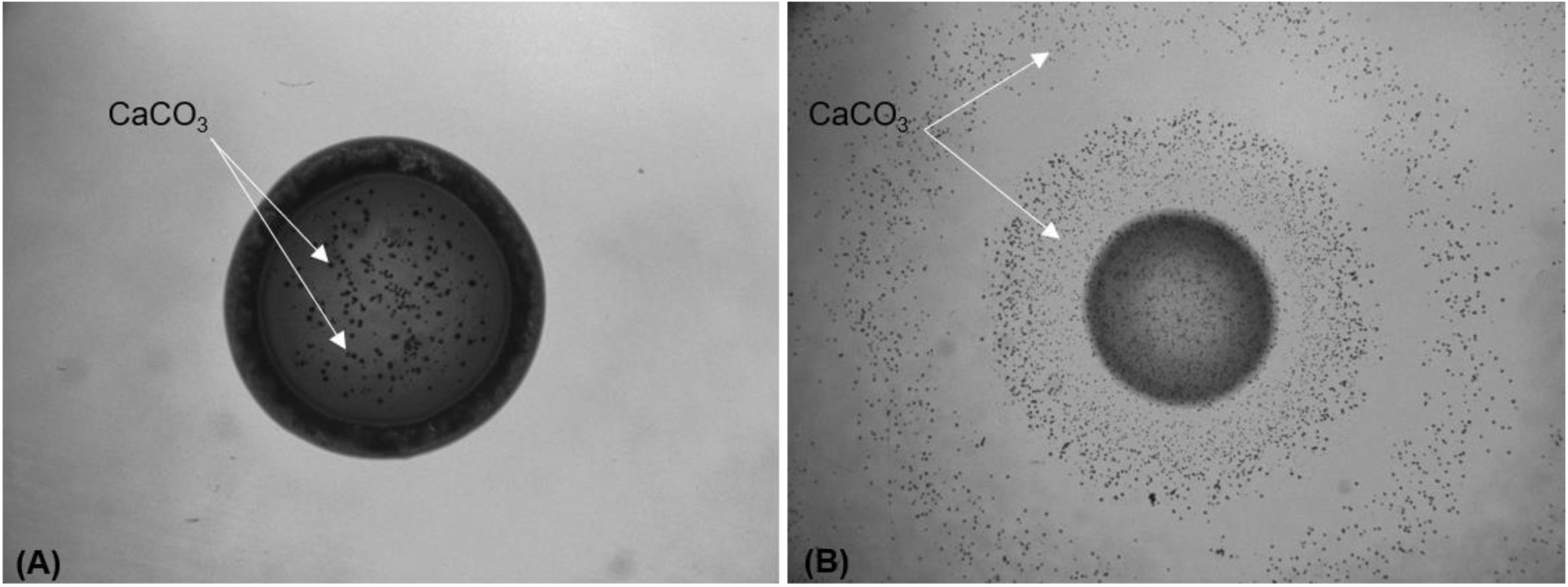
Crystal formation correlates with pH changes in non-ureolytic and ureolytic bacteria. (A) Non-ureolytic isolate PD1_1 shows localised crystals on the bacterial colony. (B) Ureolytic *S. pasteurii* demonstrates crystal precipitation throughout the media, corresponding to pH changes that occur throughout the media. Arrows indicate CaCO_3_ crystals.

Interestingly, two isolates that had tested positive for ureolysis in the test broth, CG6_1 and CG4_2, did not produce the characteristic rapid rise in pH and associated crystal precipitation when grown on solid media (Fig. 2 and S2). A possible explanation for this discrepancy might be that in these isolates, urease activity is inducible in response to a cue that was present in test broth, but not in the solid media. The main difference between the two media was the content of nitrogen, which was very low in urease test broth (0.01 % (w/v) yeast extract, 2 % (w/v) urea), but high in the solid media (0.2 % (w/v) yeast extract, 0.5 % (w/v) peptone, 1 % (w/v) Lab-Lemco powder, 2 % (w/v) urea). Nitrogen metabolism and urease production are tightly controlled in bacteria (Mobley and Hausinger, 1989). One mechanism of regulation of urease is by repression of activity in the presence of ammonia- or nitrogen-rich compounds and derepression of synthesis when nitrogen levels are low (Mobley and Hausinger, 1989). We therefore tested to see whether nitrogen levels in the media were having an effect on urease activity in these strains. Isolate CG6_1 was grown in the presence of urea in media with either low (0.02 % yeast extract, 0.0125 % peptone, 2 % urea) or high (0.2 % yeast extract, 0.125 % peptone, 2 % urea) nitrogen content, and urease activity was quantified (Fig. 4A). Urease activity in CG6_1 was high when grown in low nitrogen conditions, whilst significantly lower levels were detected in high nitrogen conditions. This suggests that urease is only active in CG6_1 when the nitrogen supply is limited. This is clearly a distinct type of behaviour from that seen in CG7_3 and *S. pasteurii*, which both utilise urea rapidly regardless of the nitrogen composition of the media (Fig. 2). However, while the overall urease activity of CG7_3 appeared similar to that of *S. pasteurii* on solid media (Fig. 2), this isolate can grow well in the absence of urea (Fig. S3 and S4), while *S. pasteurii* cannot. This suggests that CG7_3 may also be able to control the expression of its urease genes, with urea acting directly as an inducer of urease activity. We therefore tested the activity of whole cells grown in low and high nitrogen conditions as before, but also compared cells grown in the presence and absence of urea (Fig. 4B). When CG7_3 was grown in the absence of urea, it displayed medium urease activity under low nitrogen conditions, which was further repressed by high nitrogen conditions. However, when cells were grown in the presence of urea, urease activity was high regardless of nitrogen conditions. Therefore, while urease activity in CG7_3 did show a partial response to nitrogen limitation, this was overcome by incubation with urea, which clearly acted as the main inducer and may be the preferred nitrogen source for this bacterium under the chosen growth conditions. Taken together, our results indicate that there is a continuum of ureolytic activity in environmental bacteria and that this, as seen above (Fig. 2-3), will have an impact on crystal formation and biomineralization by different isolates.

**Figure 4.**
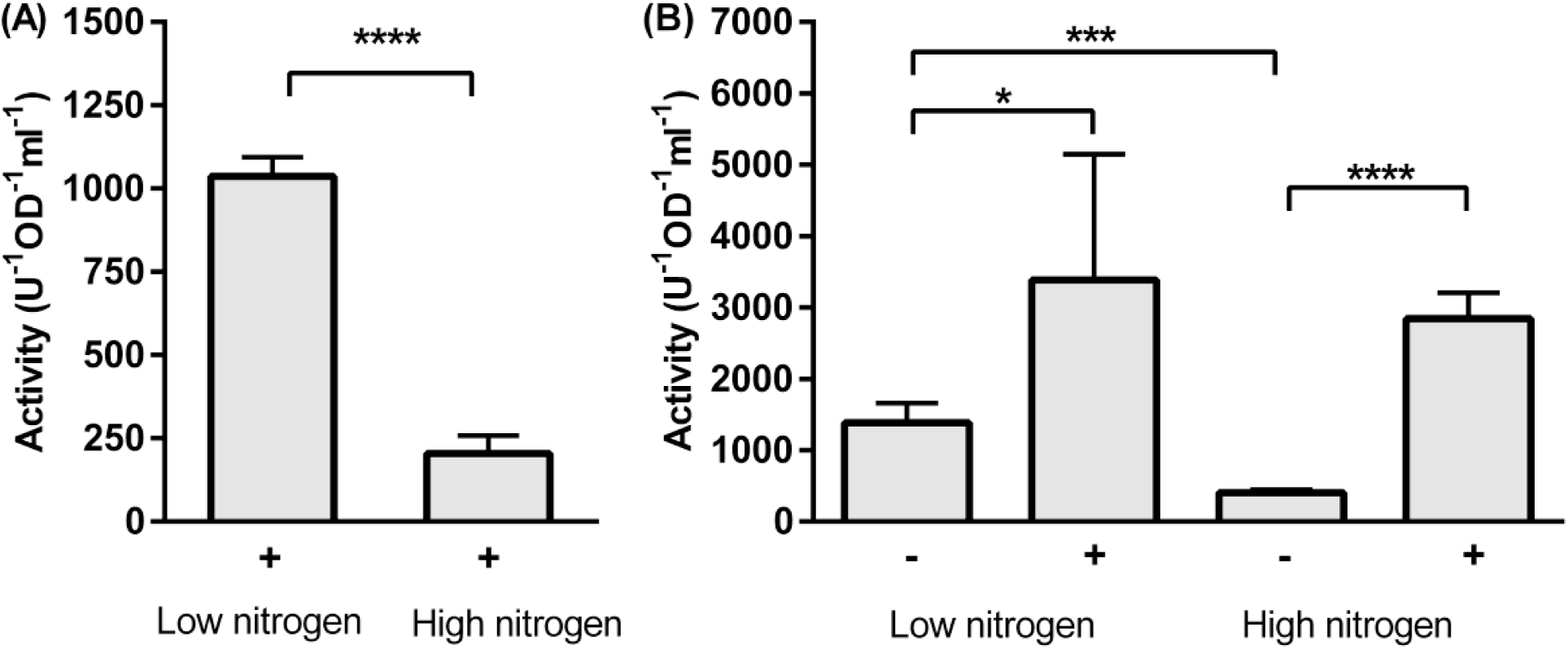
Regulation of urease activity by environmental nitrogen and urea. Isolates CG6_1 (A) and CG7_3 (B) cells were grown in high or low nitrogen conditions in the presence (+) or absence (-) of urea. Error bars represent standard deviation from 2-3 biological reps with 1-2 technical reps each. Significance was tested using an unpaired t-test (A) and one-way ANOVA followed by unpaired t-test (B). *, *P* <0.05, ***, *P* <0.001 and ****, *P* <0.0001.

### Effect of ureolysis on crystal formation

Having established that crystal formation on solid media is very different between ureolytic and non-ureolytic strains and that there are notable variations in ureolytic activity between environmental isolates, we wanted to understand how ureolysis influences crystal formation in more detail. To investigate this, we studied the kinetics of precipitation in our set of isolates. Growing isolates in liquid culture with calcium over several days allowed us to monitor the rates and total amounts of precipitate formed and correlate this to bacterial growth and global pH changes. Growth of ureolytic isolates caused rapid alkalinisation of the media, reaching pH 9 within two days, the first time point assayed. This resulted in complete precipitation of the available calcium in the media on the same time-scale, with no further change observed in two weeks of incubation (Fig. 5A and Fig. S5, *S. pasteurii*). Growth of non-ureolytic bacteria caused a more gradual increase in pH, approaching pH 9 only in the second week of incubation (Fig. 5B and Fig. S5). Precipitation of calcium in the media was also more gradual, although both ureolytic and non-ureolytic isolates attained the maximum theoretical levels of precipitation by the end of the first week. Strikingly, we observed noticeable differences in cell numbers over the experimental period. Viable cell counts decreased dramatically in ureolytic isolates and were undetectable within two days (Fig. 5A), likely due to high ammonia concentrations in the media of these isolates resulting from the cleavage of urea. Alternatively, these cells may have been encased by calcium carbonate, preventing their further growth and division. In contrast, for non-ureolytic isolates cell numbers gradually increased, before declining again over prolonged incubation (Fig. 5B). These differences in viability over time between ureolytic and non-ureolytic bacteria may have implications for application and should especially be considered in cases where the continued presence of viable cells will be required for multiple cycles of precipitation.

**Figure 5.**
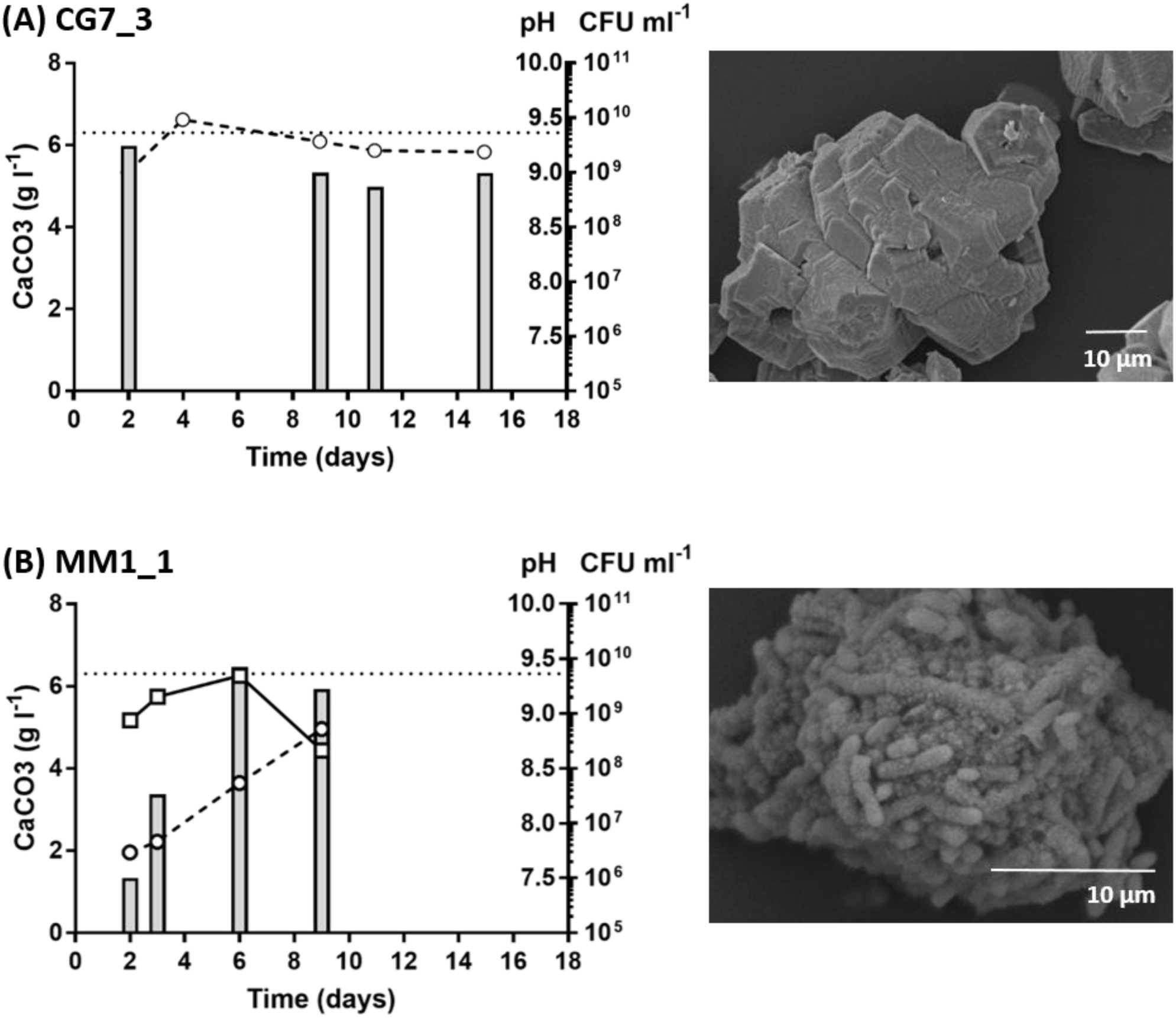
Calcite precipitation kinetics and crystal morphology in ureolytic CG7_3 and non-ureolytic MM1_1. The ureolytic strain CG7_3 (A) and the non-ureolytic MM1_1 (B) were grown in LB medium supplemented with urea (20 g l^−1^) and Ca(OAc)_2_ (10 g l^−1^) and precipitation of insoluble calcium carbonate (bars, g l^−1^), pH changes (circles) and changes in cell number (boxes, CFU ml^−1^) were monitored over time (days). Maximum theoretical level of precipitation is indicated (dotted line). *Right*, electron micrographs of representative precipitate taken at day 9.

Considering that the total amount of precipitate was similar in ureolytic and non-ureolytic isolates, we next investigated whether the differences in the speed of precipitation had any effects on the resulting crystals. As an initial observation, the precipitate recovered from ureolytic strains was a much finer powder than that recovered from non-ureolytic strains. EDX analyses of the different precipitates confirmed that all crystals tested were composed of calcium carbonate (Fig. S6). However, further investigation using SEM revealed that the crystal morphology differed between ureolytic and non-ureolytic strains (Fig. 5). When precipitation was rapid, such as in ureolytic strains CG7_3 (Fig.5) and *S. pasteurii* (Fig. S5 and Fig S7), inorganic, homogenous crystals were produced. Non-ureolytic strains such as MM1_1 that precipitated calcium carbonate more slowly produced crystals containing significant proportions of bacterial cells and appeared more “organic” in nature (Fig. 5B). Similarly, under these conditions CG4_2 and CG6_1 displayed the gradual increases in pH (Fig. S5) and more organic precipitate characteristic of non-ureolytic strains (Fig. S7). Considering that the assay conditions used for this experiment were high nitrogen, it can be expected that urease activity was repressed under these conditions and resulted in strains behaving like non-ureolytic strains. This differing appearance of precipitates in ureolytic versus non-ureolytic conditions was consistently observed across all isolates tested in this assay (Fig. S5 and Fig. S7). These results show that the differences in ureolytic activity across our environmental isolates not only affected the kinetics of biomineralization, but also had a clear impact on the crystal morphologies in the resulting precipitate.

### Profound effects of urease activity on precipitation of calcium carbonate

To exclude that the observed differences in calcium carbonate precipitation were simply due to strain-to-strain variation among our isolates, we exploited the fact that CG7_3 was capable of switching between ureolytic and non-ureolytic states depending on the availability of urea (Fig. 2 and Fig. S3). This allowed us to directly investigate the impact of ureolysis on biomineralization in a single strain. When CG7_3 was grown in the absence of urea, there was a gradual rise in pH and calcium carbonate precipitation as seen in non-ureolytic strains (Fig. 6A). In contrast, in broths supplemented with urea we observed the rapid rise in pH and associated calcium carbonate precipitation characteristic of the ureolytic strains studied above (Fig. 6B). Viable cell counts also dropped dramatically in CG7_3 utilising urea (Fig. 6B), whereas in the absence of urea CG7_3 cell numbers remained relatively stable over the first 10 days and declined by 13 days, likely due to prolonged nutrient depletion. SEM analysis of the precipitates over the time-course of the study confirmed our previous observations (Fig. 5) that urease activity was correlated with the rapid precipitation of homogenous, inorganic crystals (Fig. 6B). When assessed over time, we noticed that the initial precipitate consisted of the spherical calcium carbonate, typical of the polymorph vaterite, followed by the eventual conversion into the rhombohedral morphology associated with calcite, the more stable polymorph (Fig. 6B and Fig. S8-S9). In comparison, in the absence of urea CG7_3 precipitated calcium carbonate more gradually, and these precipitates were very organic in appearance (Fig. 6A and Fig. S8-S9). Moreover, these ‘organic’ crystals were often much larger than the inorganic spheres produced in the presence of urea, likely due to aggregation via interspersed bacterial cells. As these experiments were all performed on the same bacterial isolate, our observations are a direct reflection of the effects of urease activity on calcite precipitation. Although both pathways produced similar quantities of precipitate, there were major differences in precipitate morphologies, number of viable bacteria and pH of the bulk phase. To our knowledge, this is the first systematic study into the fundamental differences in biomineralization between different environmental bacteria. Both ureolytic and non-ureolytic bacteria are currently being developed for a range of industrial applications (DeJong *et al*., 2010). The findings reported here may therefore offer a key step towards a rational design approach to choosing which mechanism of biomineralization is better suited to specific industrial applications.

**Figure 6.**
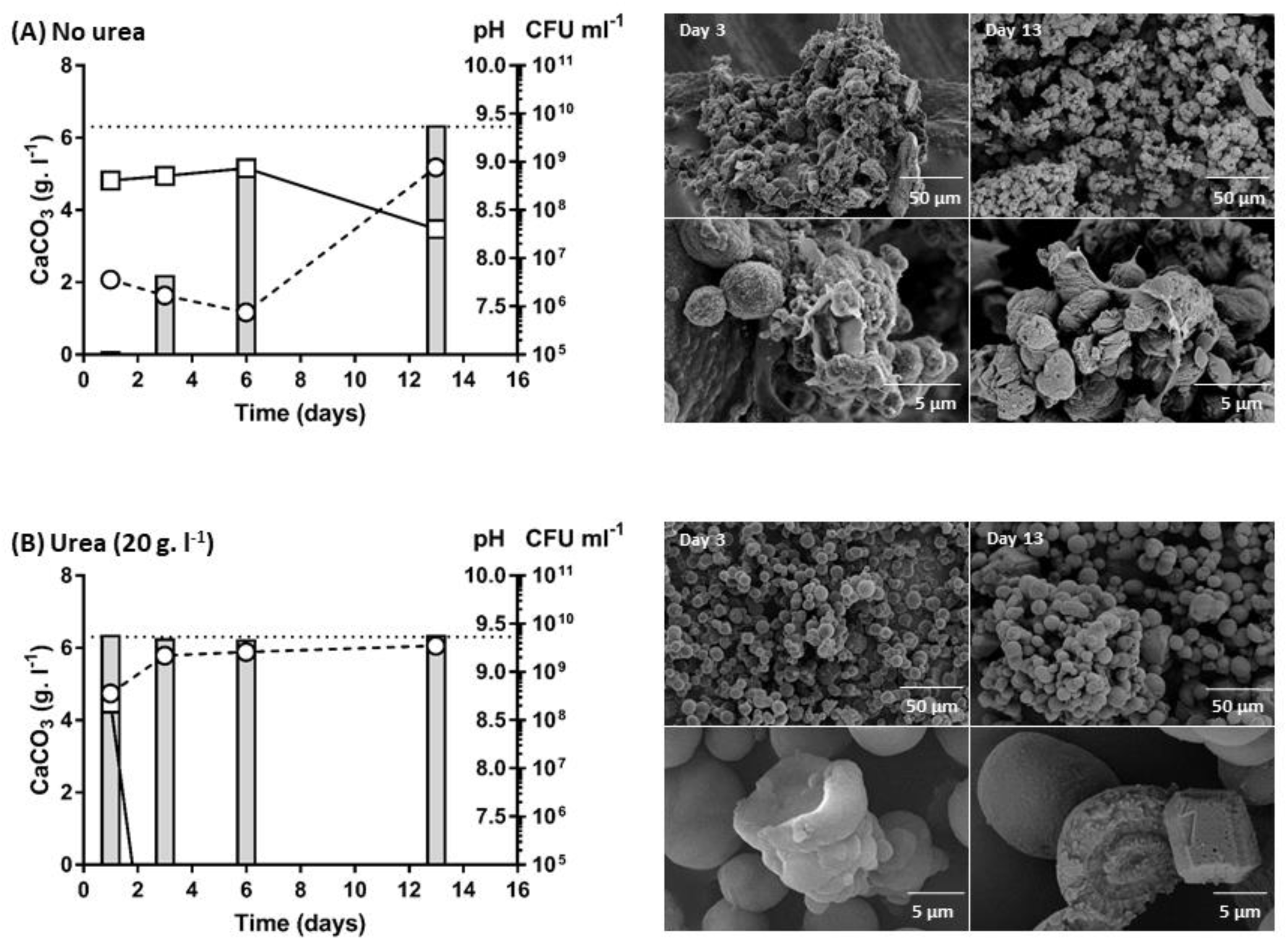
Comparison of ureolytic and non-ureolytic mechanisms of calcite precipitation in CG7_3. The ureolytic strain CG7_3 was grown in LB medium supplemented with Ca(OAc)_2_ (10 g l^−1^) in the absence (A) and presence (B) of urea (20 g l^−1^). Precipitation of insoluble calcium carbonate (bars, g l^−1^), pH changes (circles) and changes in cell number (boxes, CFU ml^−1^) were monitored over time (days). Maximum theoretical level of precipitation is indicated (dotted line). *Right*, electron micrographs of representative precipitate taken at days 3 and 13.

### Industrial relevance of different calcite precipitation strategies

To test if the differences in biomineralization mechanism translate to an applied setting, we next assessed the performance of ureolytic and non-ureolytic strains in self-healing concrete applications, using cement mortars as our test system. We produced spores for each strain, which were then encapsulated in light-weight aggregate before casting them in mortars, together with yeast extract and calcium nitrate, as well as urea in the case of ureolytic strains, to provide nutrients for bacterial growth and sufficient calcium for biomineralization. Mortars were cured and then cracked under three-point bending to obtain a target crack width of 500 µM. The self-healing process was subsequently monitored over 8 weeks.

Autogenous healing, which occurs to some degree due to cement hydration, was seen in control mortars that lacked any bacterial spores (Fig. 7 and Fig. 8). This healing was mostly observed along the top edge of the crack and rarely extended down the sides of the mortar prism. Mortars containing either ureolytic or non-ureolytic bacteria also displayed crack healing, and this generally extended across the top as well as down the sides of the crack (Fig. 7 and Fig. 8). However, healing in mortars containing ureolytic bacteria was less regular than seen with non-ureolytic strains, with the sides of cracks not consistently sealing (Fig. 8).

**Figure 7.**
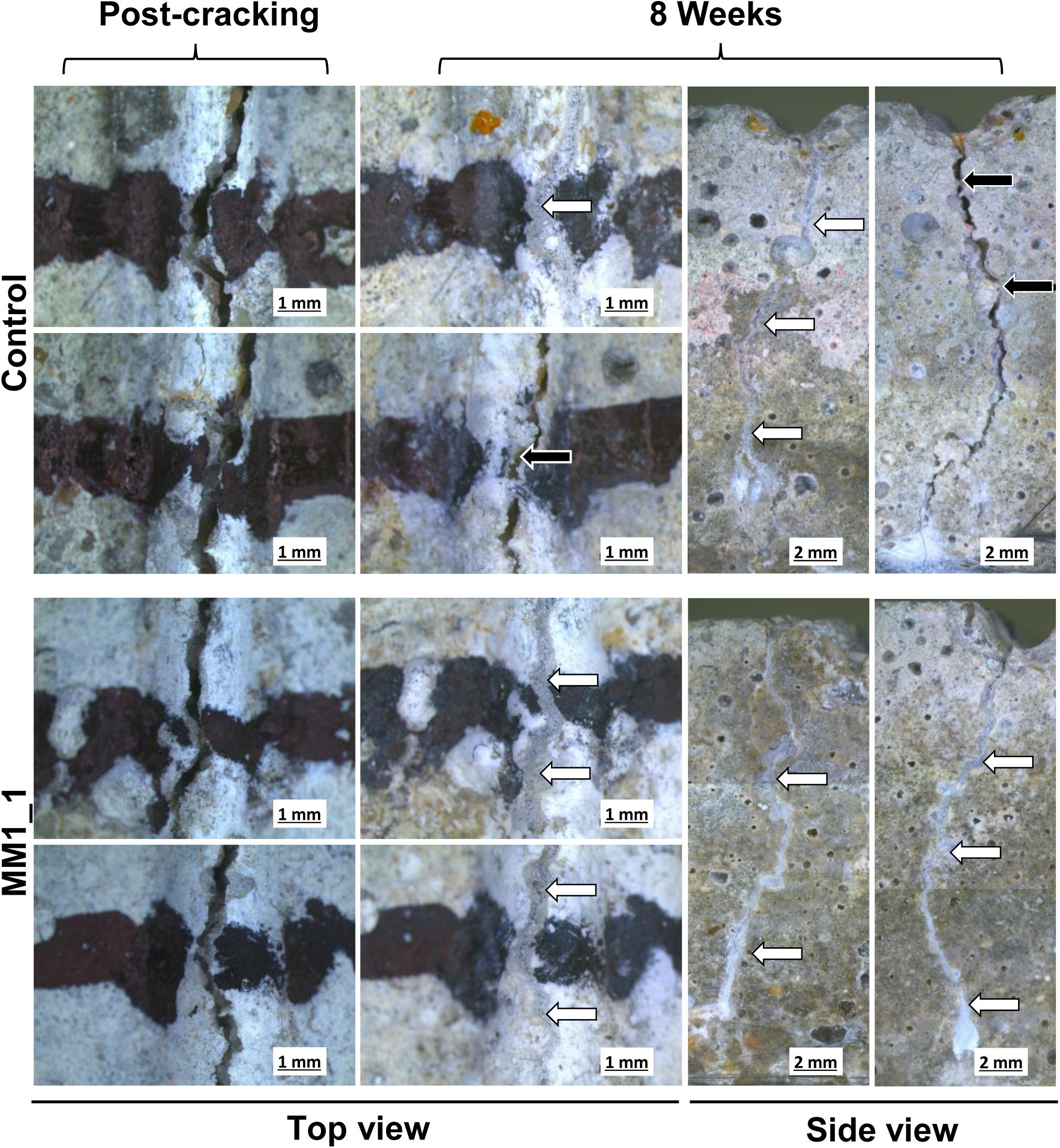
Crack closure in control mortars and mortars containing non-ureolytic bacteria. Cracks in mortar specimens are shown immediately post-cracking and after 8 weeks of healing. Cracks were marked with black pen to allow the same region to be monitored over time. Two independent regions along the top of the mortar were monitored per sample. Side views show the cracks down both sides of the mortar at 8 weeks. *White arrows*, complete crack closure. *Black arrows*, incomplete crack closure.

**Figure 8.**
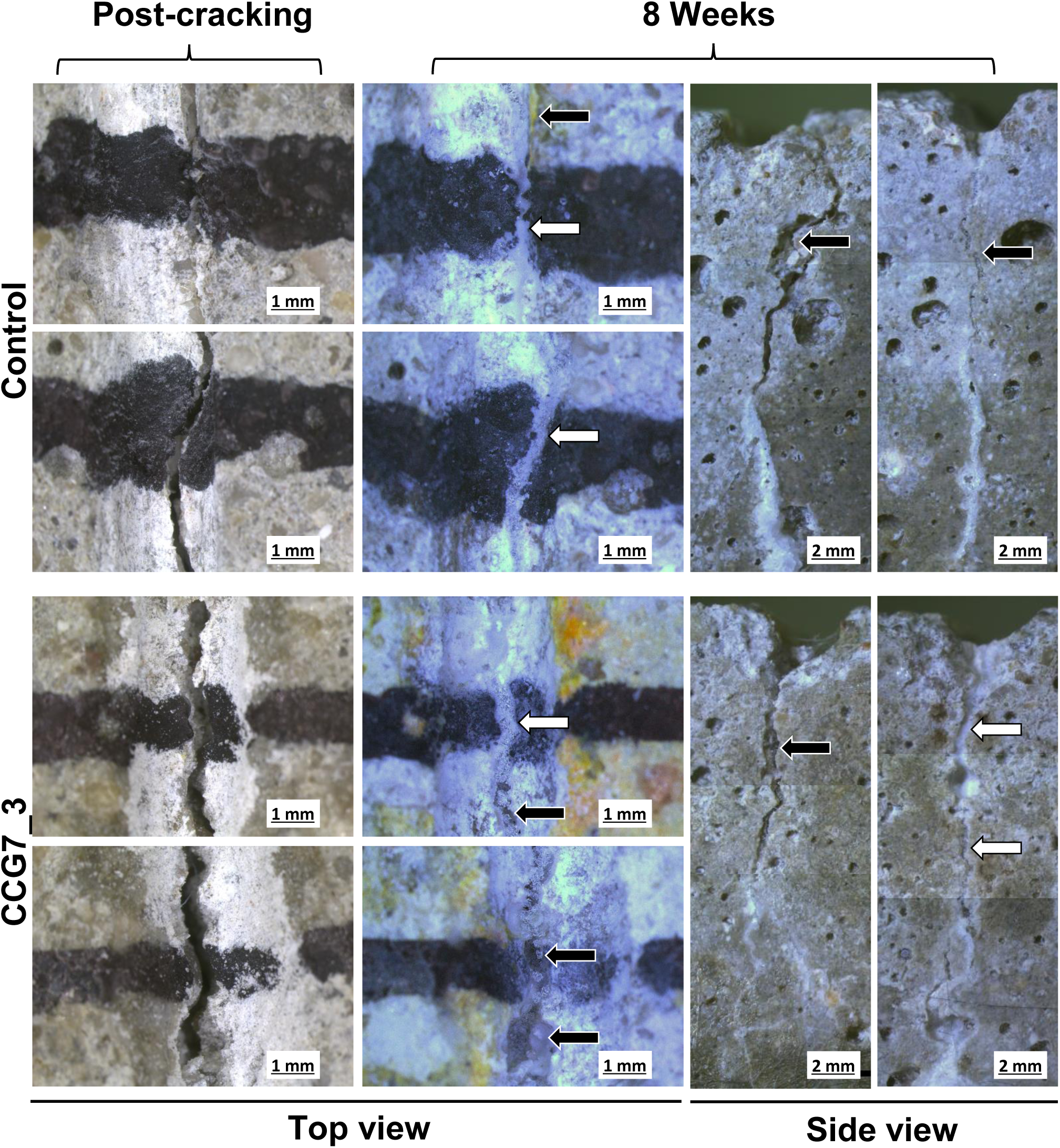
Crack closure in control mortars and mortars containing ureolytic bacteria. Cracks in mortar specimens are shown immediately post-cracking and after 8 weeks of healing. Cracks were marked with black pen to allow the same region to be monitored over time. Two independent regions along the top of the mortar were monitored per sample. Side views show the cracks down both sides of the mortar at 8 weeks. *White arrows*, complete crack closure. *Black arrows*, incomplete crack closure.

The key aim of bacteria-induced self-healing of concrete is to re-establish water tightness of the structure to prevent water ingress and subsequent corrosion of steel reinforcement. While visual inspection allowed an initial assessment of the healing process, we next sought to test water tightness of our mortars using water flow tests following eight weeks of healing. Recovery of water tightness in mortars containing only the standard cement mix, with no nutrients or additional calcium showed variable recovery of water tightness, generally close to 40 % (Fig. 9, ‘Reference’). Control mortars containing yeast extract and calcium nitrate, but without bacterial spores, showed much higher recovery (averaging 87-95 %). This recovery is likely because of a combination of autogenous healing and possible presence of environmental bacteria, which may be able to utilise the yeast extract and thus contribute to precipitation. As the mortars were not made or kept in a sterile fashion to more closely reflect industrial application, contamination with such environmental bacteria must be considered likely. The mortars containing non-ureolytic bacteria showed a strikingly consistent recovery in water tightness, leading to near complete resistance to water flow, with all specimens healing to over 90 % and many reaching close to 100% (Fig. 9). Interestingly, while mortars containing ureolytic bacteria also showed good restoration of water tightness, there was more variation in the degree of healing obtained. EM1 performed as well as the non-ureolytic strains, but CG7_3 showed lower overall recovery values (mean 87 %) that were similar to the controls lacking encapsulated bacteria. This discrepancy in results may reflect the variability in ureolytic activity in these strains. Given the clear dependence on growth conditions in some strains described above, it is difficult to predict the degree of ureolytic ability displayed in cement mortars. An alternative explanation may be that the rapid precipitation and small crystal size observed in ureolytic isolates does not reliably lead to retention of the precipitate within the crack and may thus not perform as reliably in self-healing as the larger aggregates with organic components of the non-ureolytic strains. It will be interesting to investigate the details of material performance following healing with both types of bacteria to fully understand the implications of the different mechanisms of precipitation.

**Figure 9.**
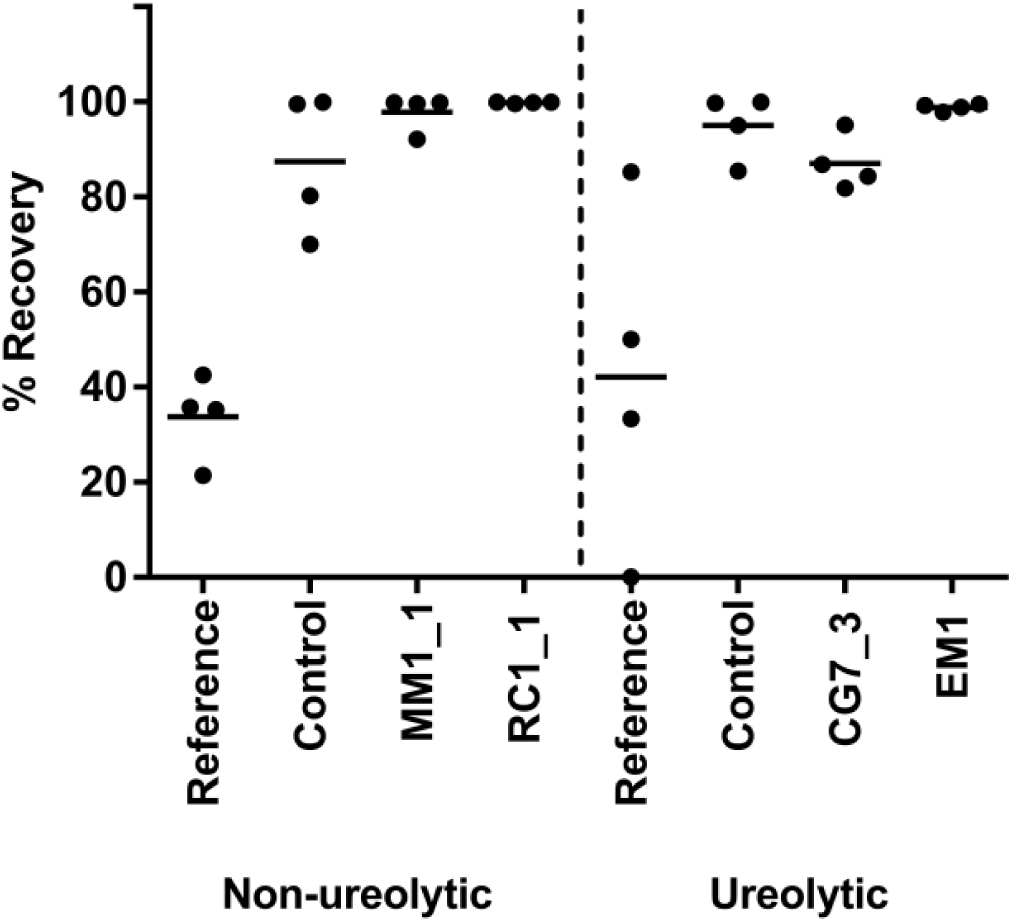
Recovery of water tightness in mortars after two months of bacteria-based self-healing. Water flow through mortars containing ureolytic (CG7_3, EM1) or non-ureolytic (MM1_1, RC1_1) bacteria was determined immediately after cracking and following 8 weeks of healing. Reference mortars contained no additives; Control mortars contained yeast extract and calcium nitrate, but no bacteria. Recovery of water tightness was determined as a percentage relative to the freshly cracked specimen (0%: no change in water flow, 100%: complete inhibition of water flow). Data is shown for four specimens per condition from two separate cracking/healing experiments.

### CONCLUDING REMARKS

The environment represents a large reservoir of potential in terms of exploiting bacteria for commercial use of MICP. There have been numerous studies on MICP for varying applications, however, most of these have been restricted to the use of a single or small number of species and were very application specific. This limits our understanding of precipitation capabilities of environmental bacteria more generally. In order to establish a stronger knowledge base to facilitate new and specialised technical applications we surveyed the ability of environmental bacteria to precipitate calcite. We found that the majority of isolates were capable of biomineralization, in line with previous results (Boquet *et al*., 1973). This is most likely due to their ability, through metabolic processes, to create a microenvironment around the cell that aids in the precipitation process. The electronegative nature of their cell surface facilitates crystal deposition on the surface (Schultze-Lam *et al*., 1996), making precipitation more favourable. Our detailed characterisation of biomineralization by our set of isolates, however, revealed fundamental differences in the way in which different bacteria precipitate calcium carbonate.

Ureolytic bacteria rapidly precipitate calcium carbonate, having led to ureolysis becoming one of the main pathways used in MICP applications to date. However, in our study, only 16% of isolates were able to cleave urea, and even amongst these there was a marked diversity in ureolytic activity. Differences in how isolates regulate urease activity were seen, with some responding to low nitrogen levels whilst others responded to urea as a primary inducer. This was in striking contrast to the constitutive high urease activity of *S. pasteuri*, one of the most frequently utilised MICP-capable bacteria to date (Stocks-Fischer *et al*., 1999;; Tobler *et al*., 2011;; Zhu *et al*., 2016). The performance of ureolytic bacteria will therefore depend on the precise conditions encountered in their environment, and this should be carefully considered when choosing the bacterial species for a given application. The importance of this point is further emphasised by our finding that ureolysis affected crystal formation both in terms of kinetics and in the morphology of the precipitate.

The majority of our isolates precipitated calcium carbonate without possessing ureolytic activity. In contrast to initial expectations that this mechanism may lead to the formation of low amounts of precipitate, we found that all of our isolates were capable of precipitating the total amount of calcium supplied in the growth media, although this process took more time with non-ureolytic strains. Importantly, we showed that the precipitate formed by the non-ureolytic bacteria consisted of larger, mixed organic/inorganic crystals. This was in good agreement with our initial observation that precipitation in these bacteria on solid media only occurred on top of the colony and therefore likely involved the physical presence of cells. The mixed nature of the precipitate means that for the same amount of calcium being used, a larger volume of precipitate can be formed. This could present a major advantage in industrial application where often only a limited amount of calcium is available to fill a relatively large space, such as in self-healing concrete.

When we tested the performance of our isolates in self-healing of cracked cement mortars, we found that, indeed, non-ureolytic bacteria caused a more consistent recovery of water tightness and more complete healing. The ureolytic bacteria tested showed less consistency in performance, although one strain gave similar results to the non-ureolytic isolates. It is difficult to ascertain whether these inconsistencies in behaviour are due to differences in ureolytic activity in the applied setting or are a consequence of the purely inorganic precipitate formed by ureolytic strains. It seems plausible that the supply of yeast extract as part of the cement mix will lead to a low-nitrogen environment conducive to urease gene expression, and the presence of urea should ensure activity in those strains that are urea-responsive. While further experimentation is needed to determine the precise conditions encountered by the bacteria within hardened cement mortars as well as the material properties of the precipitate formed, in applications where nutritional supply is hard to control, the use of non-ureolytic bacteria may give more robust and reliable performance.

Self-healing concrete of course only represents one of many potential applications of MICP and each application will have its own challenges. For example, in spray-on applications such as restoration of existing structures and historic buildings, the rapid precipitation of large amounts of precipitate through ureolysis may be advantageous. In contrast, during soil stabilisation where bacteria and their substrates are pumped into the ground, rapid rates of ureolysis could result in premature precipitation and lead to blockages at the injection site as has been reported (Cheng and Cord-Ruwisch, 2014). In this case, it would be more beneficial to use microbes that are non-ureolytic, or that switch to ureolysis at a later time point, as seen in our isolates that responded to low environmental nitrogen.

In summary, most bacteria have the ability to precipitate calcium carbonate given the right conditions, and the most suitable bacteria to use will be application dependent. While most previous studies have focussed on one or two isolates, we here show for the first time the plethora of MICP activities and capabilities the environment has to offer as a ‘talent pool’ that can offer bespoke solutions to many different applications. MICP-based technologies may offer solutions to problems caused by a rising global population and may even be used to mitigate global warming. The increasing demand for infrastructure, as well as the need to engineer land on which to build it, results in an increase in CO_2_ emissions and release of harmful chemicals into the environment (DeJong *et al*., 2010). MICP can reduce this environmental burden by producing buildings that are more durable, prolonging the life of existing buildings, and improving soil properties without the use of toxic and hazardous chemical additives. Moreover, technologies that actively remove CO_2_ from the environment and sequestrate it in harmless or even useful minerals such as calcite will be critical in meeting zero emission goals in the future.

## EXPERIMENTAL PROCEDURES

### Bacterial isolates and growth

Sample collection was from six locations across the United Kingdom including limestone caves and immature calcareous soils (Mendip Hills, England), soil, rock and limestone caves (Bath, England), soil and rock scrapings (Monmouthshire, Wales) and soil and scrapings from marine rock in two locations in Cornwall (Mount’s Bay and Falmouth). Samples were stored at 4°C until use. Of each sample, 0.5 g was re-suspended in 1 ml sterile saline solution (0.85% (w/v) NaCl) and heated to 80°C for 20 minutes to enrich for spore-formers. Selection for alkali-tolerant bacteria was carried out by plating 100 µl of this suspension onto 0.25×B4 medium (1 g l^−1^ yeast extract, 12 g l^−1^ Trizma base, 1.25 g l^−1^ glucose, 2.5 g l^−1^ calcium acetate (Ca(OAc)_2_), 15 g l^−1^ agar) adjusted with NaOH to pH 9. Glucose and Ca(OAc)_2_ were filter-sterilised and added after autoclaving. Individual colonies with unique colony morphology were re-streaked onto 0.25×B4 medium buffered with 75 mM CHES, 75 mM CAPS, adjusted with NaOH to pH 10. Strains capable of growth at pH 10 and with visible crystals on the colony or in the surrounding agar were selected for further characterisation. Single colonies were inoculated in lysogeny broth (LB;; 10 g l^−1^ tryptone, 5 g l^−1^ yeast extract, 10 g l^−1^ NaCl) and grown overnight, before storage at −80 °C in a 25% (w/v) glycerol solution. Isolates were subsequently maintained on LB agar.

*Sporosarcina pasteurii* DSM33 was included as a reference organism and maintained on LB agar supplemented with 20 g l^−1^ urea (filter-sterilized, added after autoclaving). All bacterial cultures were grown at 30 °C, and liquid cultures were agitated at 150 rpm. Growth of liquid cultures was monitored spectrophotometrically by optical density at 600 nm wavelength (OD_600_) in cuvettes of 1 cm light path length. Representative isolates showing the best characteristics were deposited into the DSMZ collection (Deutsche Sammlung von Mikroorganismen und Zellkulturen GmbH, Germany) as follows*: DSM XX (RC1_1), DSM XX (PD1_1), DSM XX (EM1), DSM XX (CG7_3), DSM XX (CG7_2), DSM XX (CG6_1), DSM XX (CG4_2), and DSM XX (MM1_1).

*\*Note to reviewers: The DSM numbers have not yet been issued but will be available by the time the manuscript has been through the review process*.

### Identification of isolates and phylogenetic analysis

The 16S rDNA gene fragment was amplified using the 27F (5’-AGAGTTTGATCMTGGCTCAG-3’) and 1492R (5’-TACCTTGTTACGACTT-3’) primers (Lane, 1991). Amplification was performed in a total volume of 25 µl containing 12.5 µl of 2x OneTaq Mastermix (New England Biolabs, NEB), 9.9 µl of nuclease-free H_2_O, 2.4 % (v/v) DMSO, 0.2 µM of each primer and 1 µl of an overnight culture as template. Reaction conditions were as follows: initial denaturation at 94 °C for 5 min followed by 30 cycles consisting of denaturation at 94°C for 30 sec, annealing at 45° C for 30 sec, and extension at 68 °C for 2 min. Final extension was at 68°C for 5 min. For enzymatic clean-up, 2 µl of EXO-SAP (100 U shrimp alkaline phosphatase (Thermo Scientific), 100 U Exonuclease I (Thermo Scientific), and 895 µl nuclease-free H_2_O) was added to every 5 µl of PCR product and the reaction was incubated at 37°C for 15 min before enzyme inactivation at 80°C for 15 min. Sequencing of PCR products was carried out using the chain termination method (Eurofins Genomics, Germany). In the case of strains where direct sequencing of the PCR product was unsuccessful, 16S rDNA fragments were first cloned into the Topo^®^ vector according to manufacturer’s instructions (TOPO TA Cloning^®^ Kit, Invitrogen). Sequencing of the resulting plasmids was performed using the M13 forward primer (5’-CCCAGTCACGACGTTGTAAAACG-3’). Bioedit (Hall, 1999) was used to assemble sequences obtained from forward and reverse sequencing reactions. Sequences were compared against the non-redundant GenBank nucleotide collection using BLASTN (http://www.ncbi.nlm.nih.gov). For phylogenetic analyses, the 16S rDNA sequences of nearest relatives for each strain according to BLASTN analysis, as well as of *Sporosarcina pasteurii* DSM 33 were obtained from GenBank, and evolutionary analyses were carried out in MEGA 7.0 (Kumar *et al*., 2016). Sequences were aligned using MUSCLE, and conserved blocks were selected from these multiple alignments using GBlocks (Castresana, 2000). Phylogenetic trees were constructed using the maximum likelihood (ML) method based on the Tamura-Nei model (Tamura and Nei, 1993). Bootstrap values were inferred from 1000 replicates and partitions reproduced in less than 50 % bootstrap replicates were collapsed.

### Analysis of calcium carbonate precipitation

Spatial distribution of calcite precipitation and pH changes was assessed by spotting 20 µl of an overnight culture (OD_600_ adjusted to 1) onto nutrient agar (Oxoid, 23 g l^−1^) supplemented with phenol red (0.025 g l^−1^) and Ca(OAc)_2_ (2.5 g l^−1^ or 10 g l^−1^), with or without urea (20 g l^−1^). Physical appearance of plates and crystals were recorded photographically at 1 hr, 3 hr, 5 hr, 24 hr, and 48 hr.

Rate of mineral precipitation over time was determined by inoculating (1:1000) 150 ml LB broth supplemented with 10 g l^−1^ Ca(OAc)_2_ from overnight cultures and monitoring viable cell numbers, pH, and insoluble calcium precipitated over time. Viable cells were determined as colony-forming units (CFU) by the plate count method. pH was recorded from aliquots of the culture using a pH electrode (Jenway 924 030, Cole-Parmer, Staffordshire UK) coupled to a pH meter (Jenway 3510 pH Meter). Insoluble precipitate was recovered by centrifugation (975 × *g*, 2 minutes at room temperature (RT)) and washed three times in 50 ml distilled water to remove planktonic cells and culture medium before oven drying at 50 °C for 48 hours. Morphology of dried precipitates was examined at accelerating voltages ranging from 10 to 20 kV by scanning electron microscopy (SEM;; JSM 6480LV, JEOL, Welwyn, UK) equipped with an energy dispersive x-ray (EDX) analyser for elemental analysis. Representative samples were prepared by spreading dried precipitate on double-sided carbon tape placed on aluminium stubs. For EDX, samples were placed under vacuum overnight before analysis. For imaging, samples were gold-coated by sputtering for 3 min at RT. Samples imaged with field emission electron microscopy (FE-SEM;; JSM-6301F cold field emission SEM, JEOL) were prepared in a similar manner as for SEM but coated in Chromium to a thickness of 20 nM and examined at 5 kV.

### Urease activity

Qualitative urease activity of bacterial isolates was determined using urease test broth (0.1 g l^−1^ yeast extract, 20 g l^−1^ urea, 0.01 g l^−1^ phenol red, 0.67 mM KH_2_PO_4_, 0.67 mM Na_2_HPO_4_ (pH 6.8±0.2). Test broth (2 ml) was inoculated with 100 µl of an overnight culture grown in LB and incubated at 30 °C for up to 5 days. A urease-positive reaction was characterised by a change in colour from yellow/orange to pink.

For quantitative urease measurements in whole cells, cultures were grown at 30 °C with agitation (150 rpm). Overnight cultures (20 µl, LB broth) were used to inoculate 25 ml (1:500) low nitrogen (0.2 g l^−1^ yeast extract, 0.125 g l^−1^ peptone, 2.5 g l^−1^ NaCl) and standard media (2 g l^−1^ yeast extract, 1.25 g l^−1^ peptone, 2.5 g l^−1^ NaCl) broths. Where pre-induction by urea was required, broths were supplemented with urea (20 g l^−1^). Cultures were grown (for 24 hr) and cells were harvested from 1 ml of culture by centrifugation (8000 *x g*, 2 min). Cells were resuspended in 1 volume 0.1 M potassium phosphate buffer (pH 8.0), and OD_600_ was recorded. Depending on the urease activities of individual strains, dilutions of cells were made to ensure the activity was within the linear range of the assay. These dilutions were recorded and accounted for during OD normalisation. Urease activity was determined according to the phenol-hypochlorite method (Natarajan, 1995) and adapted from (Achal *et al*., 2009) for whole cells. The reaction mix volumes were multiplied by the number of time-points in the assay to allow one larger volume mix to be prepared. Per time point, individual reactions contained 75 µl cell suspension, 300 µl 0.1 M potassium phosphate buffer (pH 8.0) and 750 µl 0.1 M urea solution. The mixture was incubated at 30°C and 1 ml aliquots were removed at 0, 30, 60, 90 and 120 min. At each time point, the reaction was stopped by adding 300 µl phenol nitroprusside and 300 µl alkaline hypochlorite. Final colour was developed at 37 °C for 25 min and absorbance was measured at 625 nm. Ammonium chloride (25-2500 µM) solutions were used to produce a standard curve (Fig. S1). One unit of activity was defined as the release of 1 µM of ammonia per minute, per OD (U OD^−1^ ml^−1^).

### Endospore production

Endospores of bacterial strains were prepared by inoculating 150 ml LB broth (1:1000) from an overnight culture grown at 30°C with shaking (150 rpm). This culture was again grown overnight and then cells were harvested by centrifugation (3200 × *g*, 10 min at RT) and resuspended in 750 ml Difco sporulation medium (DSM;; 8 g l^−1^ nutrient broth (Oxoid), 13.41 mM KCl, 0.49 mM MgSO_4_, adjusted to pH 7.6 with NaOH. Prior to use 1mM Ca(NO_3_)_2_, 0.01 mM MnCl_2_, and 1µM FeSO_4_ were added from a filter-sterilised stock solution (Sonenshein *et al*., 1974). Cultures were grown with agitation (150 rpm) at 30 °C for at least 48 hours before sporulation was assessed using phase contrast microscopy. When the majority of cells contained phase-bright endospores, cells were pelleted by centrifugation (3200 *× g*, 10 min at RT) and washed thrice in 10 mM Tris-HCl (pH 9) followed by 30 min treatment with chlorohexidine digluconate (0.3 mg ml^−1^) to kill vegetative cells. Washing was repeated as before and spore pellets were snap-frozen in liquid nitrogen and freeze-dried under vacuum overnight.

### Preparation of mortar samples

Mortar prisms (65 mm × 40 mm × 40 mm) were comprised of two layers. The first contained (per 3 prisms): 253.3 g sand conforming to BS EN 196-1, 92 g Portland limestone cement (CEM II A-L 32.5R), 46 g water, 1 g yeast extract, 4.55 g calcium nitrate, 3.54 g aerated concrete granules. Mortars containing ureolytic bacteria also contained urea (3.68 g per 3 prisms). Spores (2.1 × 10^10^ cfu) were resuspended in 1 ml dH_2_O and imbibed into the aerated concrete granules before drying and sealing with polyvinyl acetate (PVA, 30% (w/w)). After approximately 3 hours, the second, top layer was cast. The second layer contained standard cement mortar (Per 3 prisms: 276g standard sand, 92 g cement, 46 g water). Reference specimens were cast in two layers but only contained standard cement mortar. Specimens remained at room temperature for 24-48 hours before demoulding and subsequent curing for 28 days submersed in tap water. After curing, specimens were oven dried at 50 °C for 24 hours. The top third of the prism was wrapped with carbon fibre reinforced polymer strips to enable generation of a crack of controlled width. A notch (1.5 mm deep) was sawn at mid-span to serve as a crack initiation point. Specimens were cracked by three-point bending using a 30 kN Instron static testing frame. A crack mouth opening displacement (CMOD) gauge was used to measurement crack width. Load was applied to maintain a crack growth of 25 µm per minute, and loading was stopped when the crack width was predicted to be 500 µm wide after load removal. A marker pen was used to indicate specific crack sections to enable monitoring of the crack at the same site. Following cracking, prisms were placed in tanks, open to the atmosphere and filled with tap water to 1 cm below the top of the mortars, and were incubated at room temperature for two months. Visualisation of crack healing was monitored using a Leica M205C light microscope, and images were taken of freshly cracked mortars and after 1, 4, and 8 weeks of healing.

### Water permeability tests

Water flow rate before cracking, after cracking, and after 8 weeks of healing was determined to monitor effects of cracking and healing on water permeability in mortars as described previously (Lee and Ryou, 2016). The instrument used was based on RILEM test method II.4 (RILEM, 1987). The bottom of a 10 ml measuring cylinder (0.2 ml graduation) was removed and a 40 mm PVC pipe with 15 mm depth was fixed to the bottom. This was sealed onto the mortar with glue. This cylinder was filled with water and the time it took to drop from initial height *h*_1_ to a final height *h*_2_ was recorded. The total height was either 78.5 mm or, in the case of samples with low permeability, the height to which the water level had dropped after 30 min. The permeability coefficient was calculated according to Equation 3 (Lee and Ryou, 2016).

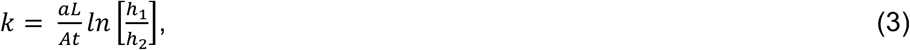

where *k* = water permeability coefficient (cm/s);; *a* = cross-sectional area of the cylinder (1.77 cm^2^);; *L* = thickness of specimen (4 cm);; *A* = cross-sectional area of the specimen which equals the cross-sectional area of acrylic plate (10.18 cm^2^);; *t* = time (s);; *h*_1_ = initial water head (12.4 cm);; *h*_2_ = final water head (cm). The percentage of crack healing was calculated as follows:

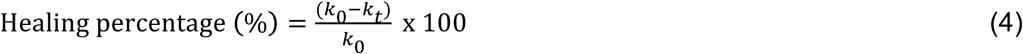

where *k*_0_ = initial water permeability after cracking, *k*_t_ = water permeability at healing time t.

## ACKNOWLEDGEMENTS

The authors would like to acknowledge the Engineering and Physical Sciences Research Council (EPSRC;; EP/PO2081X/1) and industrial collaborators/partners for funding the Resilient Materials for Life (RM4L) project. TDH was supported by a University of Bath Research Studentship Award. We thank technical staff in the Department of Architecture and Civil Engineering and the Department of Biology and Biochemistry for key support. We further thank Dr Philip Fletcher and Diane Lednitzky of the Material and Chemical Characterisation Facility (MC^2^) at University of Bath (https://doi.org/10.15125/mx6j-3r54) for their assistance with microscopy.

## FIGURE LEGENDS

**Figure S1.**
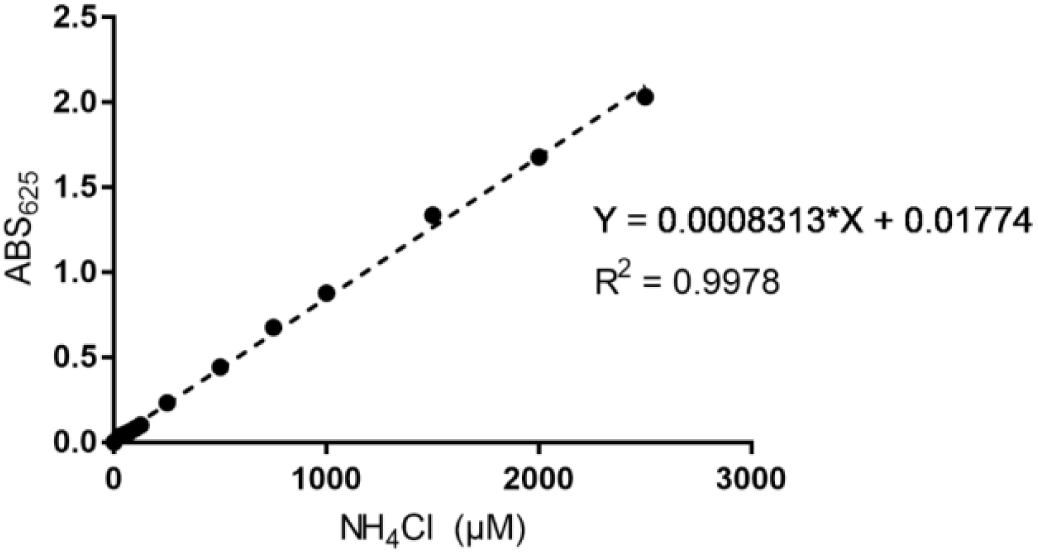
Ammonium chloride concentration vs absorbance standard curve. Ammonium chloride (25-2500 µM) solutions were used as standards to quantify the change in absorbance at 625 nm wavelength (A_625_) observed using phenol-hypochlorite as described in methods. The linear fit of ammonium chloride concentration versus absorbance used to calculate unknown concentrations is shown by the equation and dashed line.

**Figure S2.**
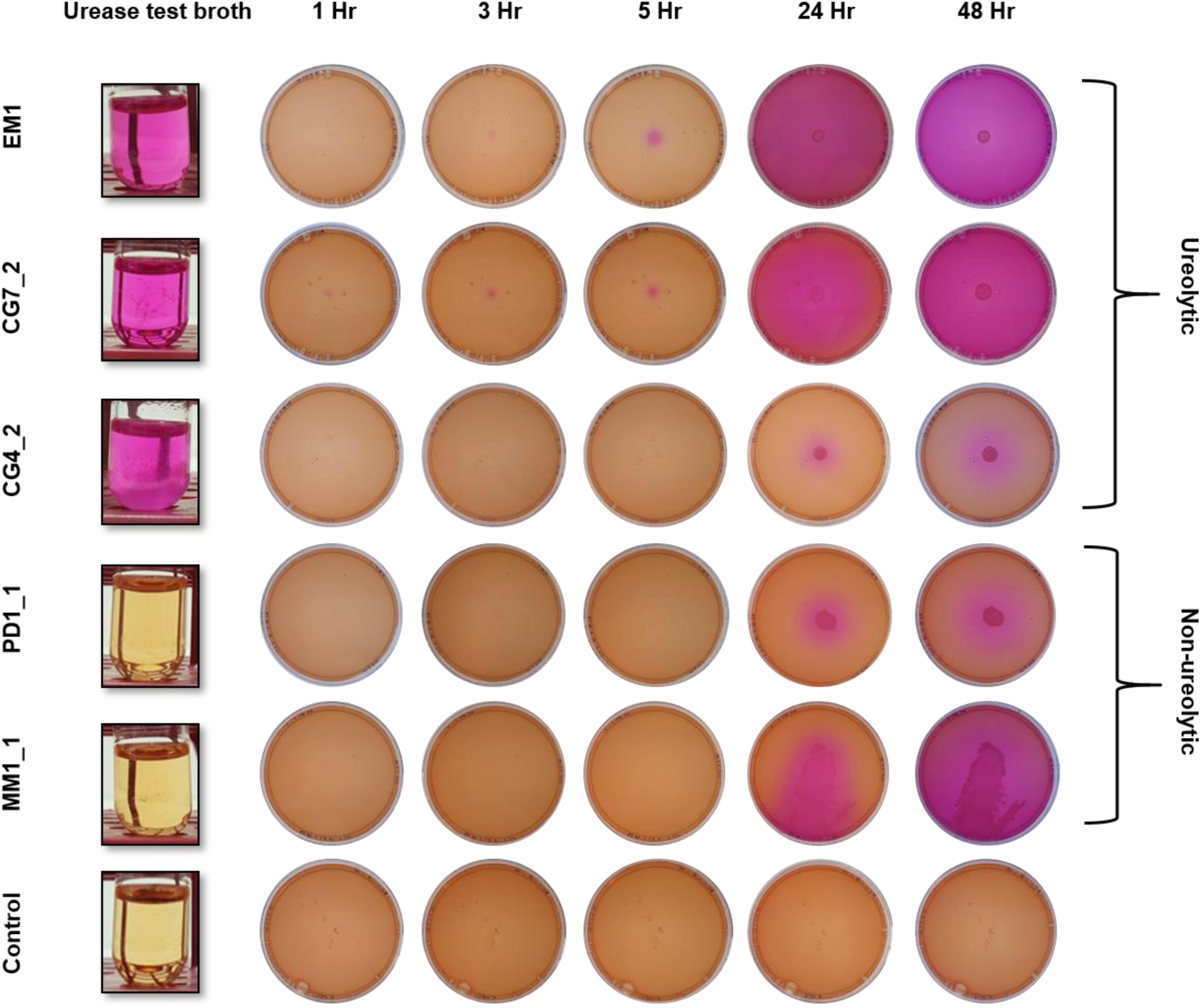
Diversity in ureolytic activity across environmental isolates. *Left*, urease test broth inoculated with the strains indicated to the left. *Right*, nutrient agar plates supplemented with phenol red, calcium acetate, and urea were inoculated in the centre with suspensions of the same strains. Pink shading indicates the pH change caused by ammonia release from ureolytic activity. MM1_1, although non-ureolytic, caused changes in pH likely due to a stress response to urea which resulted in swarming.

**Figure S3.**
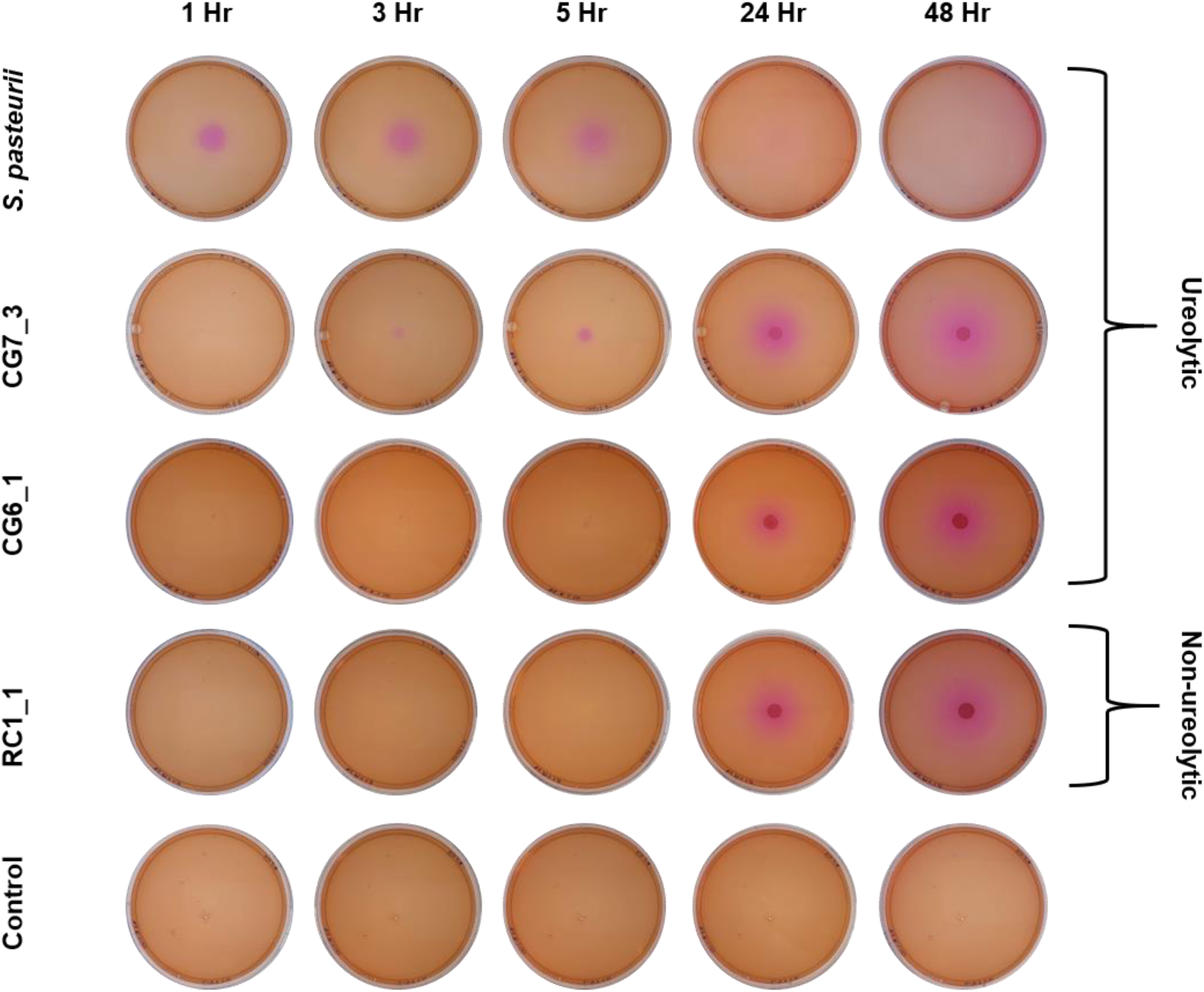
No pH changes are seen in ureolytic isolates when urea is excluded from the media. Nutrient agar plates supplemented with phenol red and calcium acetate were inoculated in the centre with suspensions of the same strains. Only local pH changes characteristic of normal metabolism are seen.

**Figure S4.**
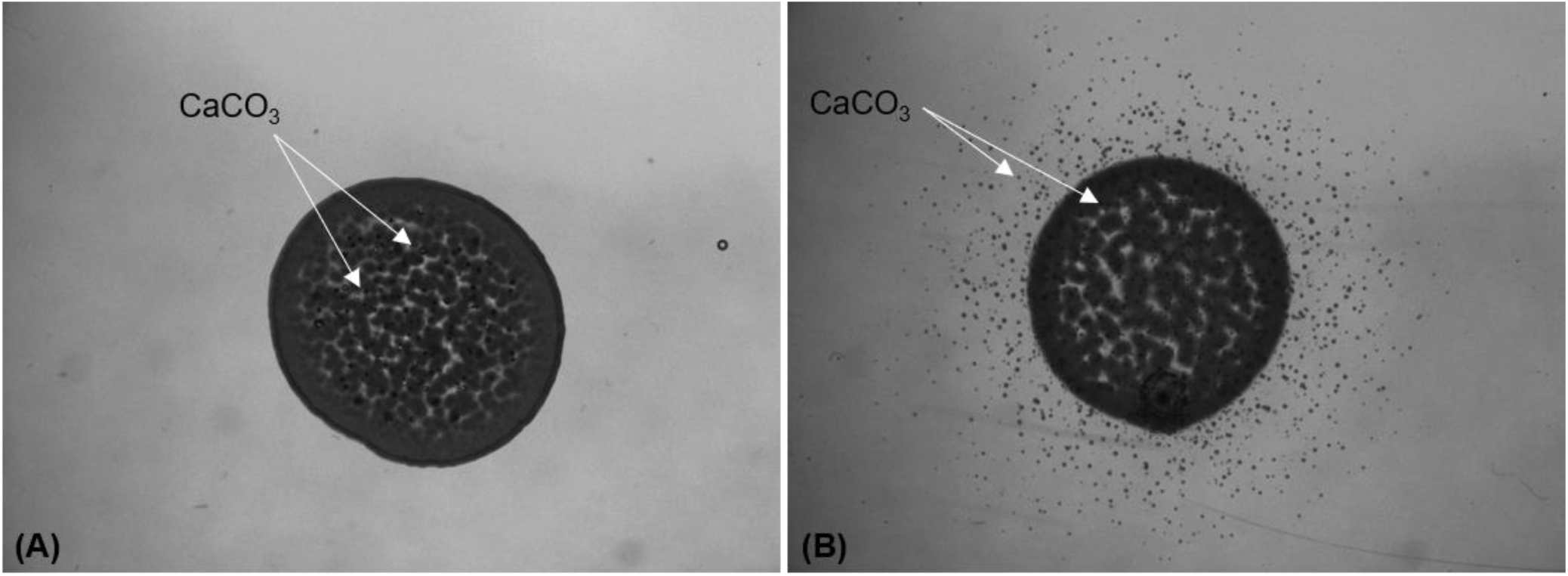
Comparison of crystal localisation in ureolytic strain CG7_3 without (A) and with (B) urea. Ureolytic bacteria precipitate crystals locally in the absence of urea, similar to what is seen in non-ureolytic strains. Arrows indicate CaCO_3_ crystals.

**Figure S5.**
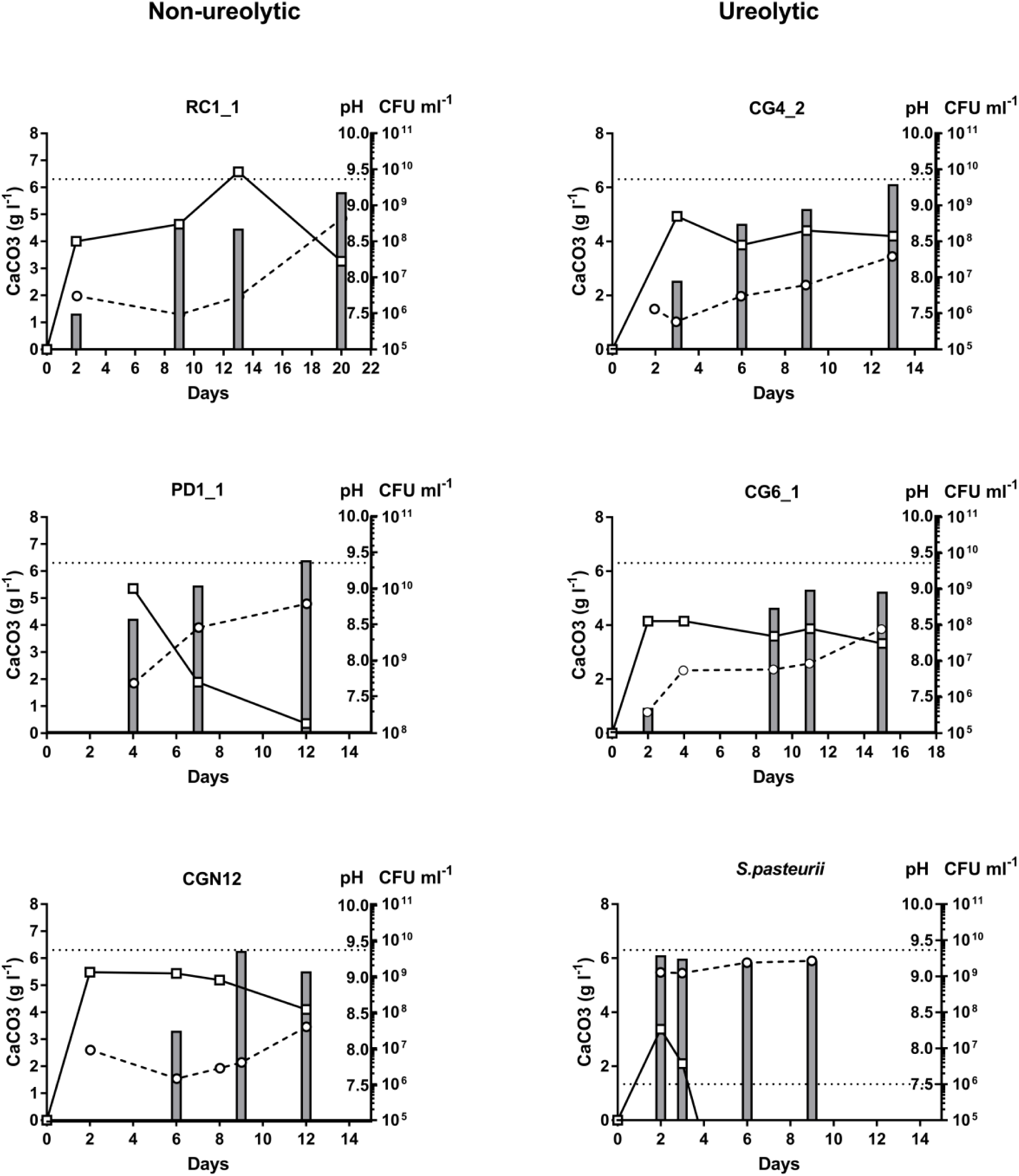
Calcite precipitation kinetics in ureolytic and non-ureolytic isolates. Isolates were grown in LB medium supplemented with urea (20 g l^−1^) and CaAoc (10 g l^−1^) and precipitation of insoluble calcium carbonate (bars, g l^−1^), pH changes (circles) and changes in cell number (boxes, CFU ml^−1^) were monitored over time (days).

**Figure S6.**
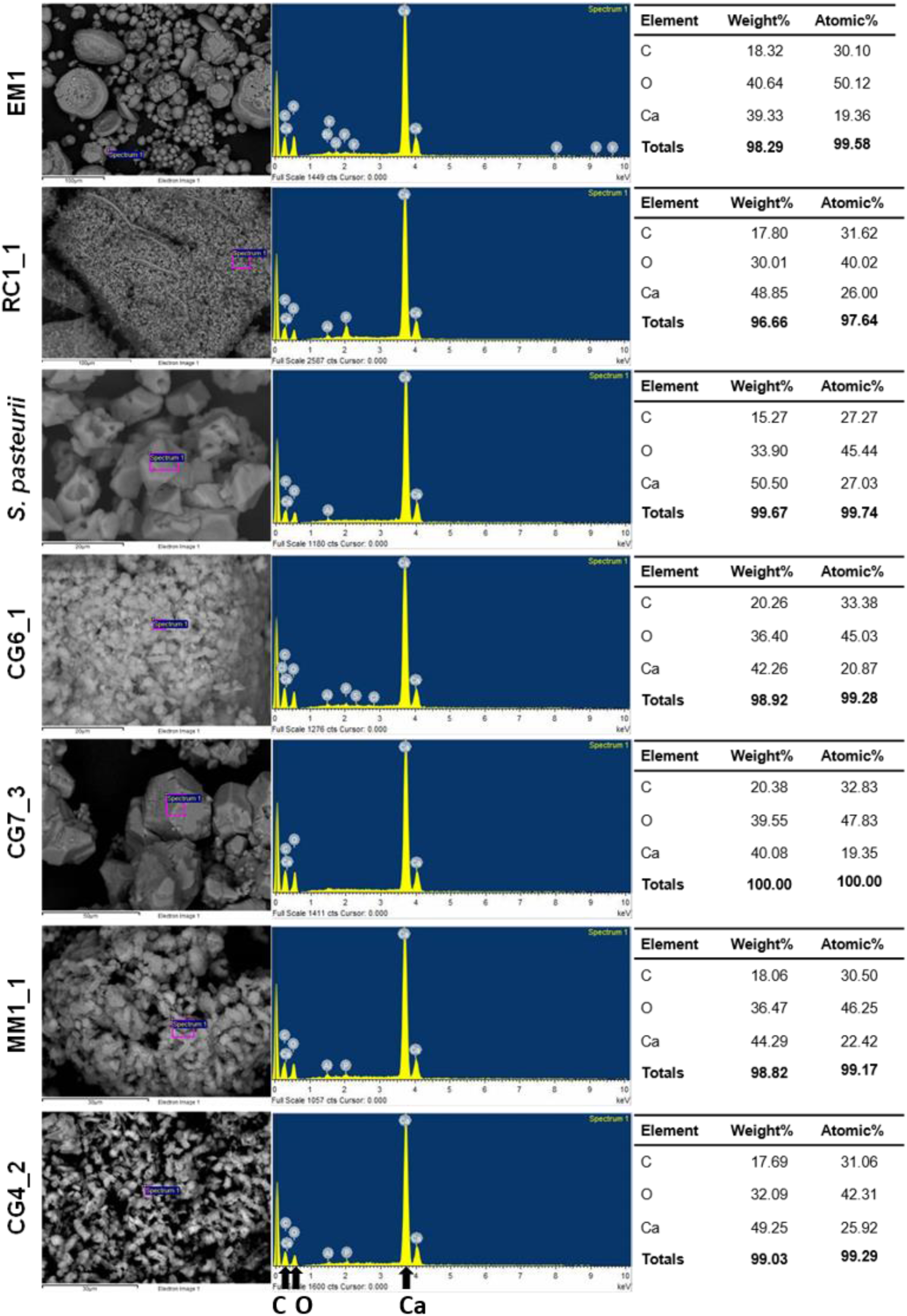
EDX analyses of precipitates produced by environmental bacteria. *Left*, the position on the precipitate where EDX was conducted (purple boxes). *Centre*, corresponding EDX spectra are shown, with the three peaks of main components labelled below. *Right*, elemental analysis indicating that the three main components (%) of the precipitate where carbon (C), oxygen (O), and calcium (Ca). Minor elements were excluded from the summary tables.

**Figure S7.**
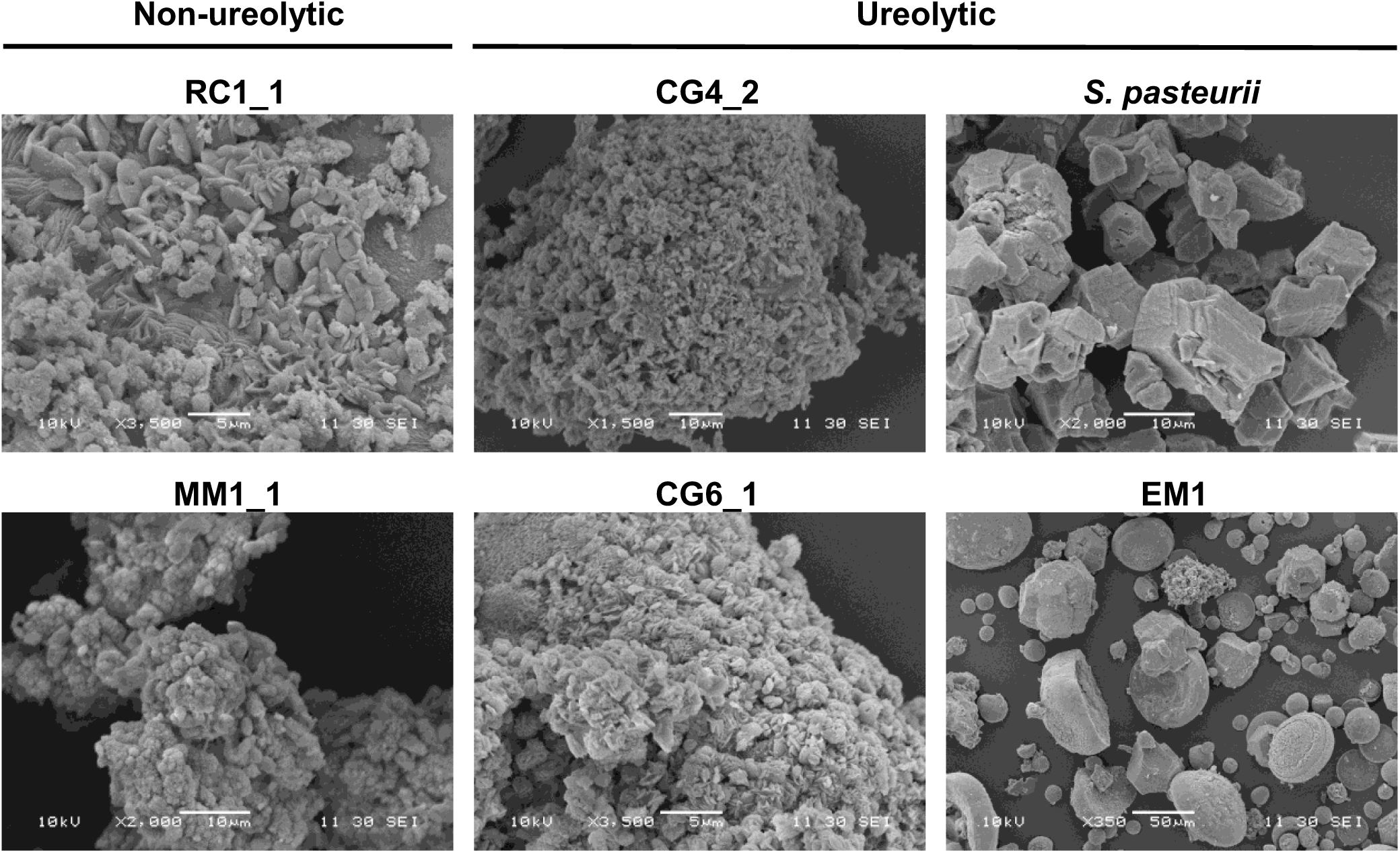
Crystal morphology in ureolytic and non-ureolytic isolates. Isolates were grown in LB medium supplemented with urea (20 g l^−1^) and Ca(OAc)_2_ (10 g l^−1^) and electron micrographs of representative precipitates were taken between days 9-15. Strain names and their ureolytic potential are stated above.

**Figure S8.**
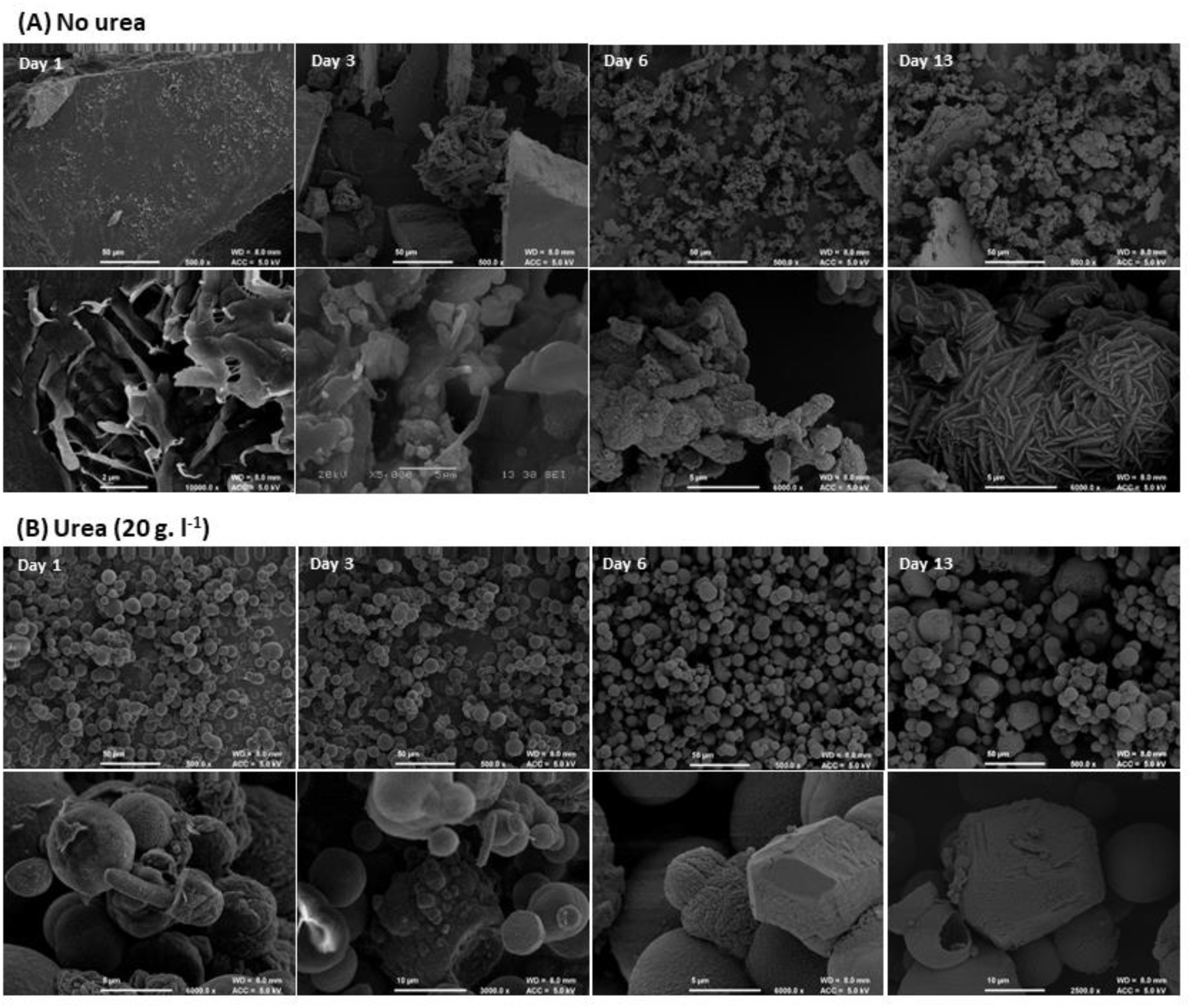
Comparison of crystal morphology in CG7_3 grown in the absence and presence of urea. The ureolytic strain CG7_3 was grown in LB medium supplemented with Ca(OAc)_2_ (10 g l^−1^) in the absence (A) and presence (B) of urea (20 g l^−1^). Electron micrographs of precipitate were taken at days 1,3, 6 and 13. The top panel of each series represents a low magnification view, the bottom panel a high magnification close-up.

**Figure S9.**
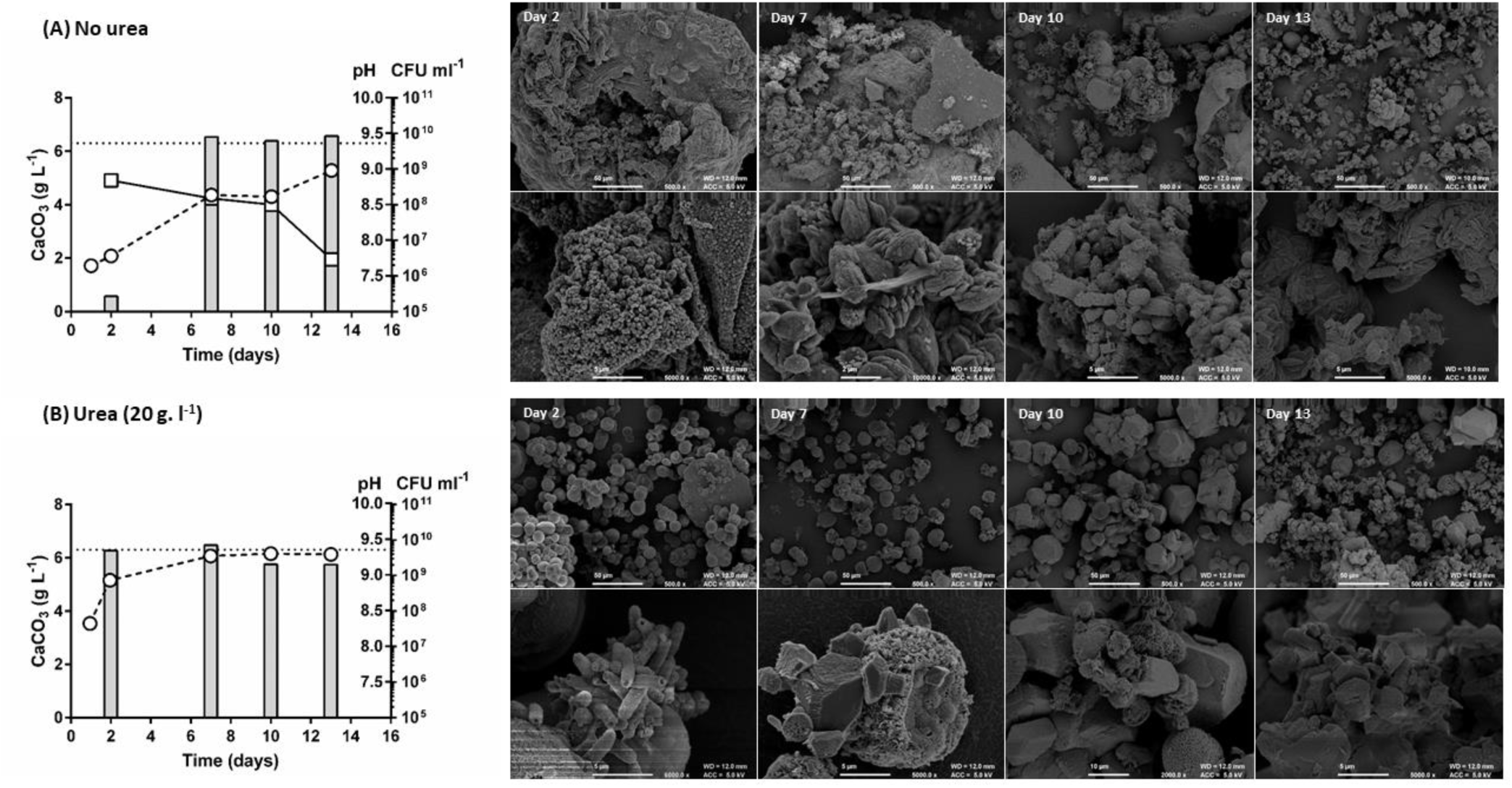
Comparison of ureolytic and non-ureolytic mechanisms of calcite precipitation in CG7_3. Ureolytic strain CG7_3 was grown in LB medium supplemented with Ca(OAc)_2_ (10 g l^−1^) in the absence (A) and presence (B) of urea (20 g l^−1^). Precipitation of insoluble calcium carbonate (bars, g l^−1^), pH changes (circles) and changes in cell number (boxes, CFU ml^−1^) were monitored over time (days). *Right*, electron micrographs of representative precipitate taken at days 2, 7, 10 and 13.

